# Inactivation of the Hippo Tumor Suppressor Pathway Promotes Melanoma

**DOI:** 10.1101/2021.05.04.442615

**Authors:** Marc A. Vittoria, Nathan Kingston, Eric Xia, Rui Hong, Lee Huang, Shayna McDonald, Andrew Tilston-Lunel, Revati Darp, Joshua Campbell, Deborah Lang, Xiaowei Xu, Craig Ceol, Xaralabos Varelas, Neil J. Ganem

**Author notes:** Correspondence to: Neil J. Ganem, Neil J. Ganem, The Cancer Center, Boston University School of Medicine, 72 E. Concord St. K-712C, Boston, MA 02118.

## Abstract

Human melanomas are commonly driven by activating mutations in *BRAF*, which promote melanocyte proliferation through constitutive stimulation of the MAPK pathway. However, oncogenic *BRAF* alone is insufficient to promote melanoma; instead, its expression merely induces a transient burst of proliferation that ultimately ceases with the development of benign nevi (*i*.*e*. moles) comprised of growth-arrested melanocytes. The tumor suppressive mechanisms that induce this melanocytic growth arrest remain poorly understood. Recent modeling studies have suggested that the growth arrest of nevus melanocytes is not solely due to oncogene activation in individual cells, but rather due to cells sensing and responding to their collective overgrowth, similar to what occurs in normal tissues. This cell growth arrest is reminiscent of the arrest induced by activation of the Hippo tumor suppressor pathway, which is an evolutionarily conserved pathway known to regulate organ size. Herein, we demonstrate that oncogenic BRAF signaling activates the Hippo pathway *in vitro*, which leads to inhibition of the pro-growth transcriptional co-activators YAP and TAZ, ultimately promoting the growth arrest of melanocytes. We also provide evidence that the Hippo tumor suppressor pathway is activated in growth-arrested nevus melanocytes *in vivo*, both from single-cell sequencing of mouse models of nevogenesis and human tissue samples. Mechanistically, we observe that oncogenic BRAF promotes both ERK-dependent alterations in the actin cytoskeleton and whole-genome-doubling events, and that these two effects independently promote Hippo pathway activation. Lastly, we demonstrate that abrogation of the Hippo pathway, via melanocyte-specific deletion of the Hippo kinases *Lats1/2*, enables oncogenic *BRAF*-expressing melanocytes to bypass nevus formation, thus leading to the rapid onset of melanoma with 100% penetrance. This model is clinically relevant, as co-heterozygous loss of *LATS1/2* is observed in ∼15% of human melanomas. Collectively, our data reveal that the Hippo pathway enforces the stable growth arrest of nevus melanocytes and therefore represents a critical and previously unappreciated barrier to melanoma development.

## Introduction

Cutaneous melanoma arises from the malignant transformation of melanocytes, which are neural crest-derived cells mainly localized to the basal layer of the epidermis. When locally resected, melanoma is highly curable; however, melanoma is the most aggressive of all skin cancers and distant-stage disease is associated with significant lethality (1). Unraveling the molecular features underlying the pathogenesis of cutaneous melanoma is essential for the development of preventative and therapeutic treatment strategies.

The vast majority of melanocytic neoplasms are initiated by oncogenic mutations in the mitogen-activated protein kinase (MAPK) pathway, with activating mutations in *BRAF* and *NRAS* occurring in ∼50% and ∼20% of cutaneous melanomas, respectively (2). Within *BRAF*-mutant melanomas, the most common activating mutation results from a single amino acid substitution from a valine to a glutamic acid generating the constitutively active mutant *BRAF*^*V600E*^ (3, 4). Despite strongly inducing proliferative signaling, melanocyte-specific expression of *BRAF*^*V600E*^ is insufficient to induce melanoma in multiple animal models; instead, *BRAF*^*V600E*^ expression leads to the development of benign nevi (moles) comprised of growth-arrested melanocytes (5-10). This is corroborated by clinical evidence as melanocytes within benign human nevi also frequently contain *BRAF*^*V600E*^ mutations (11, 12) and these melanocytic nevi rarely transform into melanoma (annual rate < 0.0005%) (13). Similarly, mutations within *NRAS* are commonly detected in congenital nevi and oncogenic *NRAS* expression in melanocytes *in vivo* does not rapidly yield melanoma (14-16). Although the risk of any single melanocytic nevus transforming into melanoma is minimal, understanding how such transformations occur is paramount as roughly one third of all melanomas co-exist with or arise from nevi (17).

These observations indicate that tumor suppression mechanisms restrain melanoma development following the acquisition of activating MAPK pathway mutations in melanocytes. A longstanding view is that strong oncogenic signals driven by mutations in MAPK pathway components leads to oncogene-induced cellular senescence (OIS), which safeguards against tumorigenesis (18-20). Supporting this view, it has been demonstrated that expression of *BRAF*^*V600E*^ in primary melanocytes *in vitro* induces an immediate cell cycle arrest and that these arrested melanocytes exhibit all of the hallmarks of oncogene-induced senescence: they become large, flat, vacuolar, express p16^INK4A^, display senescence-associated β-galactosidase (SA-β-gal) activity, and have increased heterochromatic foci and DNA damage (18, 21).

However, while it is clear that oncogene-induced senescence occurs *in vitro*, the extent to which this mechanism operates to ward off tumorigenesis *in vivo* remains unclear (22-24). Several pieces of evidence argue against OIS as being the predominant mechanism restraining the proliferation of melanocytes harboring oncogenic mutations *in vivo* (extensively reviewed in (16)). Most notably, oncogene expression (e.g. *BRAF*^*V600E*^) in melanocytes does not induce an immediate proliferative block *in vivo*. Rather, these oncogenes initially induce proliferation, as evidenced by the clonal outgrowth of melanocytes that ultimately form a nevus, which requires many rounds of cell division. Furthermore, melanocytes lacking proteins known to enforce senescence, such as p16 and p53, retain the capacity to enter a growth-arrested state, as melanocytes in *Braf*^*V600E*^*/Cdkn2a*^*-/-*^ and *Braf*^*V600E*^*/Trp53*^*-/-*^ mouse models still primarily form nevi, with only a rare few melanocytes stochastically transforming into melanoma (7).

Collectively, these data suggest that additional tumor suppressive mechanisms have the capacity to restrain the proliferation of *Braf*^*V600E*^-positive mouse melanocytes, independent of inducing senescence. Recent modeling studies have led to the postulation that the growth arrest of nevus melanocytes is not solely due to oncogene activation and OIS in individual cells, but rather due to cells sensing and responding to their collective overgrowth, similar to what occurs in normal tissues (25). This cell growth arrest is very reminiscent of the arrest induced by activation of the Hippo tumor suppressor pathway, which is an evolutionarily conserved pathway known to regulate organ size. When the Hippo pathway is activated, the Hippo kinases LATS1/2 phosphorylate the transcriptional co-activators YAP (*YAP1*) and TAZ (*WWTR1*), resulting in their inactivation by nuclear exclusion and subsequent degradation (26, 27). In contrast, when the Hippo pathway is inactivated, YAP and TAZ are active and form DNA-binding complexes with the TEAD family of transcription factors, which act synergistically with AP-1 complexes to stimulate the expression of genes mediating entry into S-phase and cell proliferation (28, 29).

It is not known if Hippo pathway activation contributes to the growth arrest of nevus melanocytes. Moreover, while Hippo pathway inactivation has been suggested to promote cutaneous melanoma growth and invasion (30-32), it remains unknown whether Hippo inactivation is sufficient to induce cutaneous melanoma initiation and/or progression. Here, we use a combination of *in vitro* and *in vivo* model systems to examine the role of the Hippo tumor suppressor pathway in restraining melanoma development.

## Results

### *BRAF*^*V600E*^ expression activates the Hippo tumor suppressor pathway *in vitro*

We sought to examine if expression of *BRAF*^*V600E*^ is sufficient to induce activation of the Hippo tumor suppressor pathway in cultured melanocytes. Previous studies using primary melanocytes have demonstrated that exogenous expression of oncogenic *BRAF*^*V600E*^ leads to an immediate p53-dependent growth arrest (9, 33). We therefore developed a system in which we could induce *BRAF*^*V600E*^ expression without an immediate cell cycle arrest in an attempt to explore Hippo pathway activation over multiple cell cycles. To do so, we generated a doxycycline-inducible system to permit controlled *BRAF*^*V600E*^ expression in non-transformed Simian Virus 40 (SV-40) immortalized melanocytes (Mel-ST cells) (34). Expression of the SV-40 early region, which encodes the small and large T viral antigens, imparts immortality to primary melanocytes via multiple mechanisms including impairment of the p53/Rb pathways (34). Induction of *BRAF*^*V600E*^ expression in Mel-ST cells increased the phosphorylation levels of the downstream kinases ERK and RSK, indicating that the cell model successfully hyperactivates MAPK signaling upon addition of doxycycline (dox) (Figure 1A).

**Figure 1:**
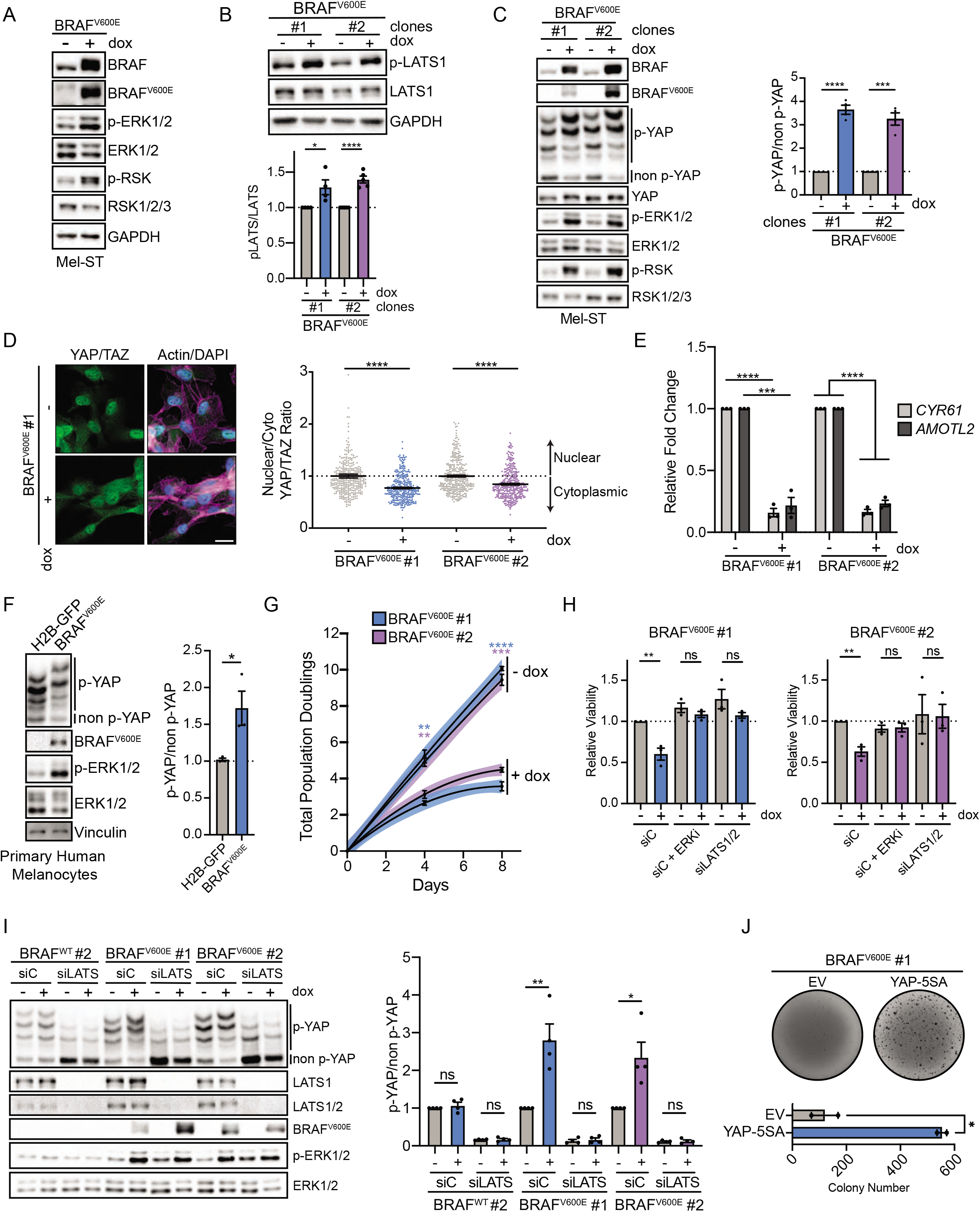
*BRAF*^*V600E*^ Activates the Hippo Tumor Suppressor Pathway. **(A)** Representative immunoblot (IB) of *BRAF*^*V600E*^ dox-inducible Mel-ST cells cultured ± dox for 24 h. **(B)** IB of indicated *BRAF*^*V600E*^ dox-inducible Mel-ST clones cultured ± dox for 24 h (n ≥ 4 independent experiments, graph shows mean relative intensity ± SEM, two-tailed unpaired t test). **(C)** Left, IB of indicated *BRAF*^*V600E*^ dox-inducible Mel-ST clones cultured ± dox for 24 h; Right, intensity quantification ratio of YAP phosphorylation from phos-tag gel (n = 4 independent experiments, graph shows mean relative intensity ± SEM, two-tailed unpaired t test). **(D)** Left, representative immunofluorescence staining of YAP/TAZ or Merge of DAPI (blue), YAP/TAZ (green), and Phalloidin (magenta) in indicated *BRAF*^*V600E*^ Mel-ST clone; Right, quantification of nuclear to cytoplasmic ratio of mean YAP/TAZ fluorescence (n > 300 cells from three independent experiments, graphs show mean ± SEM, scale bar 25 µm, two-tailed Mann-Whitney test). **(E)** Relative expression of indicated genes in *BRAF*^*V600E*^ Mel-ST clones cultured ± dox for 24 h (n = 3 independent experiments, graphs show mean ± SEM, two-tailed unpaired t test). **(F)** Left, IB of primary human melanocytes infected with indicated lentiviruses; Right, intensity quantification ratio of YAP phos-tag (n = 3 independent experiments, graphs show mean ± SEM, two-tailed unpaired t test) **(G)** Population doubling assay of indicated Mel-ST cell lines over indicated time (n = 3 independent experiments, lines represent second order polynomial non-linear fit, fill color represents 95% confidence interval of said non-linear fit, dots show mean ± SEM, color of stars indicates cell line compared, two-tailed unpaired t test). **(H)** Relative viability of Mel-ST cell lines treated with indicated siRNA or drugs for a 4 day period (n = 3 independent experiments in technical quintuplicate, graphs show mean ± SEM, two-tailed unpaired t test). **(I)** Left, representative IB of indicated dox-inducible Mel-ST clones treated with control siRNA (siC) or LATS1 and LATS2 siRNAs (siLATS) ± dox for 24 h; Right, intensity quantification ratio of YAP phos-tag (n = 4 independent experiments, graphs show mean ± SEM, two-tailed unpaired t test). **(J)** Representative crystal violet stain of *BRAF*^*V600E*^ dox-inducible Mel-ST cell line expressing indicated genes grown in dox containing soft agar with quantification below (n = 2 independent experiments in technical triplicate, graph shows mean ± SEM, two-tailed unpaired t test). ns = non-significant, \**P* < 0.05, \*\**P* < 0.01, \*\*\**P* < 0.001, \*\*\*\**P* < 0.0001

To examine if *BRAF*^*V600E*^ activates the Hippo tumor suppressor pathway *in vitro*, we induced *BRAF*^*V600E*^ expression and examined the relative levels of active LATS1/2 that is phosphorylated at the hydrophobic motif (T1079) (27). We found a significant increase in LATS phosphorylation following expression of oncogenic *BRAF*^*V600E*^ (Figure 1B). We then assessed total YAP phosphorylation (p-YAP) via phos-tag gel electrophoresis. We observed that *BRAF*^*V600E*^ induction promoted phosphorylation of YAP at multiple sites (Figure 1C). Consequently, expression of *BRAF*^*V600E*^ led to nuclear exclusion of YAP and a corresponding decrease in the expression of the YAP target genes *CYR61* and *AMOTL2* (Figures 1D and E). The observed effects on LATS and YAP activity were due to *BRAF*^*V600E*^, as overexpression of wild-type *BRAF* had no effect on LATS or YAP phosphorylation (Figures S1A-D). We further confirmed these results in multiple cell lines, including non-immortalized primary adult human melanocytes with an intact p53 pathway (Figures 1F and S1E). Importantly, the observed effects of Hippo pathway activation were not limited to expression of *BRAF*^*V600E*^ alone, as we also found that inducible expression of oncogenic *NRAS*^*Q61R*^ similarly activates the Hippo pathway (Figure S1F). Collectively, these data demonstrate that hyperstimulation of the MAPK signaling pathway through expression of oncogenic *BRAF*^*V600E*^ or *NRAS*^*Q61R*^ leads to activation of the Hippo tumor suppressor pathway *in vitro*.

We noted that expression of *BRAF*^*V600E*^ reduced melanocyte cell number ∼30-40% relative to uninduced controls over a 4-day period, despite the fact that these melanocytes were SV-40 immortalized (Figure 1G). Live-cell imaging and population doubling assays revealed this was predominantly due to a proliferative arrest, rather than increased cell death (Figures 1G, 2A-B). To test whether the observed Hippo pathway activation induced by *BRAF*^*V600E*^ expression was responsible for this proliferative defect we used RNAi to knockdown the LATS1/2 kinases in the context of *BRAF*^*V600E*^ expression. We found that loss of LATS1/2 prevented YAP phosphorylation following induction of *BRAF*^*V600E*^ and fully rescued cell growth and viability (Figure 1H-I). We then validated this result using soft agar growth assays. While parental Mel-ST cells and Mel-ST cells expressing *BRAF*^*V600E*^ fail to efficiently grow under anchorage independent conditions, Mel-ST cells expressing *BRAF*^*V600E*^ together with a constitutively active YAP mutant (YAP-5SA) demonstrated significant colony formation (Figure 1J and data not shown). These data reveal that functional inactivation of the Hippo pathway, through either LATS1/2 depletion or constitutive YAP activation, is sufficient to restore proliferation to *BRAF*^*V600E*^-expressing immortalized melanocytes *in vitro*.

### BRAF^V600E^ Promotes Mitotic Slippage and Whole-Genome Doubling

We sought to understand the mechanisms through which *BRAF*^*V600E*^ activates the Hippo tumor suppressor pathway. Disruption of MAPK signaling has been shown to promote mitotic errors and previously we have demonstrated that tetraploid cells, generated by cytokinesis failure, activate the Hippo pathway (35-37). Thus, we speculated that expression of *BRAF*^*V600E*^ may lead to Hippo pathway activation by disrupting the normal completion of mitosis.

To test this possibility, we performed live-cell imaging of doxycycline-inducible *BRAF*^*V600E*^ Mel-ST cells stably expressing the chromosome marker histone 2B-GFP (H2B-GFP). We observed that upon entering mitosis, cells expressing *BRAF*^*V600E*^ often exhibited widely-oscillating chromosomes and were unable to maintain a tightly-aligned metaphase plate relative to uninduced controls (Figure 2A). These chromosome alignment defects impaired the ability of many cells to satisfy the spindle assembly checkpoint, and consequently a portion of the *BRAF*^*V600E*^-expressing cells endured a significantly prolonged mitosis (Figure 2A-B). On average, cells expressing *BRAF*^*V600E*^ experienced a mitosis that was double the length of uninduced control cells (control: ∼74 min, induced: ∼154 min).

**Figure 2:**
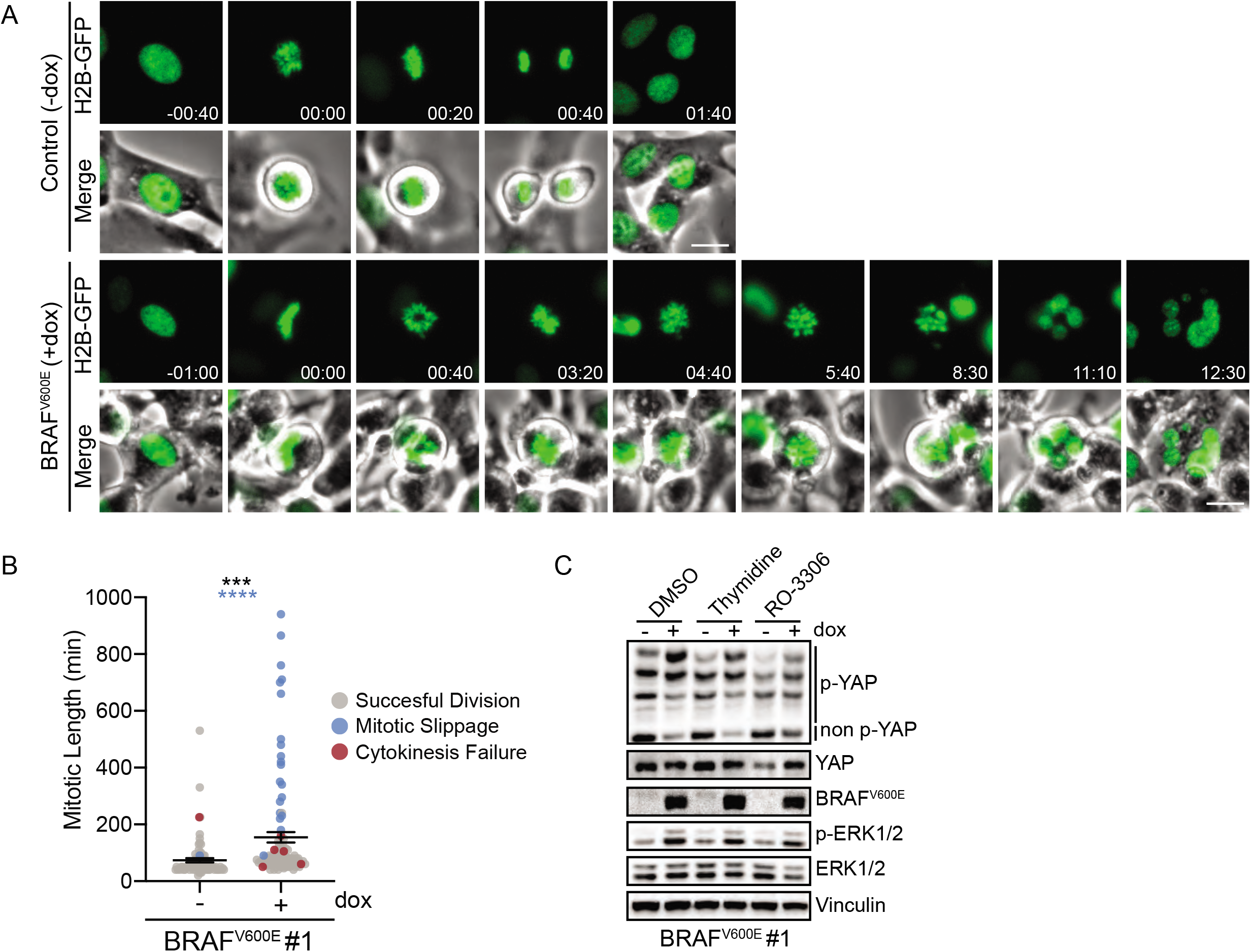
*BRAF*^*VB00E*^ Expression Induces Whole-Genome Doubling Events. **(A)** Representative still fluorescence and bright-field images from a live-cell video of H2B-GFP (green) expressing *BRAF*^*V600E*^ dox-inducible Mel-ST cells cultured ± dox (scale bar 25 μ m, hh:mm). **(B)** Plot of mitotic length and fate of individually tracked mitoses from (A) (n > 80 mitoses per condition from two independent experiments, dots represent individually tracked mitoses, black stars represent mitotic length significance, two-tailed unpaired t test, blue stars represent significance for difference in frequency of whole-genome doubling events, two-sided Fisher’s exact test). **(C)** IB of indicated Mel-ST cell line treated with indicated drugs (n = 3 independent experiments all with similar results). \*\*\**P* < 0.001, ***** < 0.0001

Cells that cannot satisfy the spindle assembly checkpoint either undergo mitotic cell death, or exit from mitosis without undergoing cell division, a phenomenon termed mitotic slippage (38). Cells that undergo mitotic slippage often generate multinucleated tetraploid cells, and multinucleated melanocytes have been observed in human nevi (37, 38). We observed that upon induction of *BRAF*^*V600E*^, the number of mitoses resulting in mitotic slippage increased significantly (control: ∼1%, induced: ∼20%) (Figure 2B). We also noted cytokinesis failure following anaphase onset was increased following *BRAF*^*V600E*^ expression (control: ∼1%, induced: ∼5%). Overall, melanocytes expressing *BRAF*^*V600E*^ experienced a significantly increased occurrence of whole-genome doubling events (control: 2.47%, induced: 24.75%). We further confirmed the generality of these findings in another independent cell line (Figures S2A-B). These data demonstrate that *BRAF*^*V600E*^, the most prevalent MAPK mutation in benign nevi, can impair mitosis leading to mitotic slippage and the formation of multinucleated tetraploid melanocytes. Indeed, this may be one mechanism by which multinucleated melanocytes form in human nevi. In support of these conclusions, it has recently been discovered that melanocyte-specific expression of *BRAF*^*V600E*^ or *NRAS*^*Q61K*^ in zebrafish leads to the formation of fish nevi largely comprised of binucleated tetraploid cells, suggesting oncogenic MAPK signaling can promote mitotic errors and tetraploidization *in vivo* as well (Darp et al, unpublished).

We examined whether *BRAF*^*V600E*^-induced tetraploidization was driving YAP phosphorylation, however our data suggested that mitotic errors leading to tetraploidization were not the major underlying cause of Hippo pathway activation in *BRAF*^*V600E*^-expressing melanocytes. First, *BRAF*^*V600E*^-expressing Mel-ST cells arrested in G_1_ (via thymidine) or G_2_ (via RO-3306 mediated CKD1 inhibition) still experienced Hippo activation despite their inability to become tetraploid (Figure 2C). Second, our immunofluorescence experiments revealed mononucleated diploid cells also exhibit decreased nuclear YAP/TAZ, demonstrating tetraploidization was not necessary to observe Hippo pathway activation (Figure 1D).

### *BRAF*^*V600E*^-induced Hippo activation is ERK-dependent and partially mediated by changes in the actin cytoskeleton

We examined whether *BRAF*^*V600E*^ specifically, or rather hyperactivation of the MAPK pathway generally, is responsible for Hippo pathway activation. We found that dampening of MAPK signaling via inhibition of the downstream kinases MEK1/2 or ERK1/2 fully prevented Hippo pathway activation, as measured by YAP/TAZ phosphorylation status, in *BRAF*^*V600E*^-expressing Mel-ST cells (Figure 3A). These data demonstrated that Hippo pathway activation is entirely mediated by hyperactivation of MAPK signaling and requires factors downstream of ERK. We further confirmed this in another independent cell line (Figure S3A). This result discounts the possibility that oncogenic *BRAF*^*V600E*^ activates Hippo signaling via direct phosphorylation of key Hippo pathway components.

**Figure 3:**
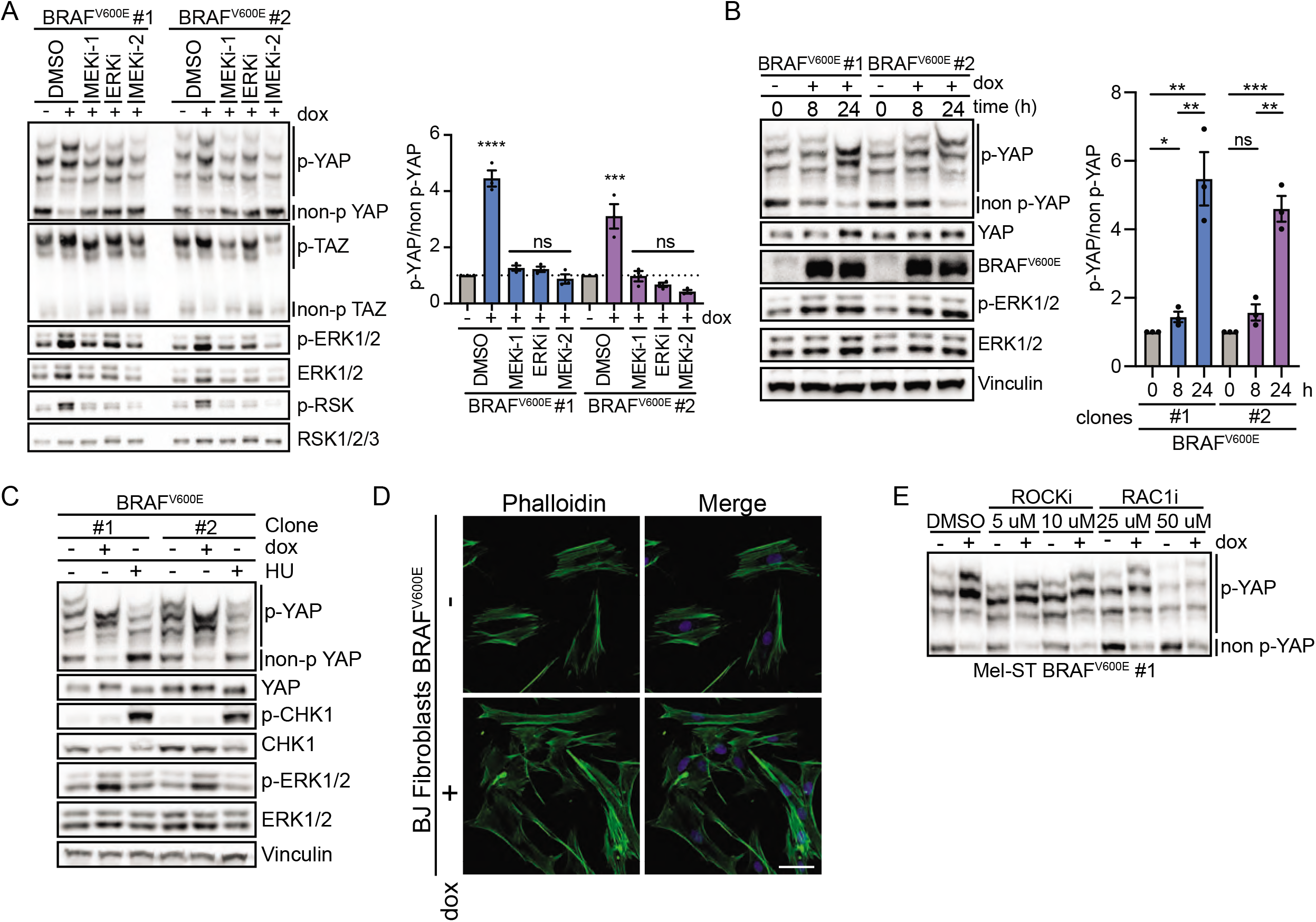
Prolonged MAPK Activation Leads to Cytoskeletal Modifications and Hippo Activation. **(A)** Left, IB of indicated *BRAF*^*V600E*^ dox-inducible Mel-ST cell lines cultured ± dox for 24 ; along with indicated drugs at the following doses: ERKi (20 nM), MEKi-1 (10 μM), MEKi-2 (20 nM); Right, intensity quantification ratio of YAP phos-tag (n = 3 independent experiments, graphs show mean ± SEM, one-way ANOVA with multiple comparisons). **(B)** Left, IB of *BRAF*^*V600E*^ dox-inducible Mel-ST cell lines cultured ± dox for indicated time; Right, intensity quantification ratio of YAP phos-tag (n = 3 independent experiments, graphs show mean ± SEM, two-tailed unpaired t test). **(C)** Representative IB of *BRAF*^*V600E*^ dox-inducible Mel-ST cell lines treated ± dox for 24 ; or with 1 mM Hydroxyurea for 6 ; (n = 3 independent experiments all with similar results). **(D)** Representative maximum intensity projections of confocal z-stacks of *BRAF*^*V600E*^ dox-inducible BJ Fibroblasts stained for phalloidin and DAPI (n = 2 independent experiments with similar results, scale bar 25 μm). **(E)** Representative IB of *BRAF*^*V600E*^ dox-inducible Mel-ST cell line treated with indicated drugs ± dox for 24 h (n = 3 independent experiments with similar results, quantification shown in Figure S4F). ns = non-significant, ** < 0.05, *** < 0.01, **** < 0.001, ***** < 0.0001

We noted that phosphorylation of YAP following *BRAF*^*V600E*^ expression requires sustained MAPK stimulation over a period of 12-16 h, as transient MAPK activation only minimally affected YAP phosphorylation (Figure 3B, S3B-C). Based on this result, we speculated that mounting oncogene-induced replication stress may be promoting Hippo pathway activation. However induction of replication stress by hydroxyurea treatment alone was not sufficient to activate the Hippo pathway (Figure 3C).

Alternatively, it has been demonstrated that oncogenic activation of the MAPK pathway dramatically alters actomyosin cytoskeletal contractility and reduces RhoA activity in an ERK1/2-dependent manner (39-41). Reductions in active RhoA are known to promote Hippo pathway activation and, furthermore, ERK1/2-dependent cytoskeletal changes have previously been shown to modulate YAP/TAZ activity in melanoma cell lines (42). We therefore posited that reduction of RhoA activity may represent one mechanism by which *BRAF*^*V600E*^-expressing cells activate the Hippo pathway *in vitro*. Indeed, we observed that there was a significant reduction in the number of actin stress fibers following activation of *BRAF*^*V600E*^, indicating reduced RhoA activity (Figure 3D, S3D-E). We also found that inhibition of Rac1, which is known to antagonize RhoA, mitigated *BRAF*-induced Hippo pathway activation (Figure 3E and S3F). These data suggest that *BRAF*^*V600E*^-induced Hippo pathway activation is at least partially mediated by prolonged MAPK hyperstimulation leading to ERK1/2-dependent cytoskeletal dysregulation. Supporting this view, endogenous *Braf*^*V600E*^ expression in mouse embryonic fibroblasts has been shown to drastically reduce actin stress fibers, and a recent study has also demonstrated expression of *BRAF*^*V600E*^ in RPE-1 cells leads to decreased RhoA activity (Darp et al. unpublished) (43).

### Growth-arrested melanocytes in benign nevi show evidence of Hippo pathway activation

It is well established that melanocyte-specific expression of *Braf*^*V600E*^ in animal models gives rise to benign nevi that harbor non-proliferating melanocytes. We hypothesized that these *Braf*^*V600E*^ -positive melanocytes may also demonstrate evidence of Hippo pathway activation, similar to our observed results *in vitro*. To test this possibility, we analyzed a single-cell RNA sequencing dataset of whole-skin extracts collected at P30 and P50 from wild-type control mice and tamoxifen-painted *Tyr::CreER*^*T2*^/*Braf*^*CA*^ mice expressing active *Braf*^*V600E*^ (25). We hypothesized any difference in Hippo activity may be only detectable at the transcriptomic level as we suspected Hippo activation *in vivo* to be subtle. We interrogated this dataset to examine if YAP/TAZ-dependent gene transcription is repressed in melanocytes expressing oncogenic *Braf*^*V600E*^. Following dimensionality reduction and initial clustering, melanocytes were identified utilizing the melanocyte-lineage marker *Dct* as previously reported (Figure 4A) (25). This cluster was also found to have high expression of another melanocyte-lineage marker, *Mlana* (Figure S4A). We then employed variance-adjusted Mahalanobis (VAM), a scRNA-seq optimized approach to obtain accurate signaling pathway scores, to examine if YAP/TAZ-mediated gene expression was decreased in *Braf*^*V600E*^*-*expressing mouse melanocytes relative to wild-type melanocytes utilizing previously published YAP/TAZ gene expression profiles (44-46). The most basic analysis, where all single-cells were binned by genotype, demonstrated that YAP/TAZ gene expression targets were significantly reduced in *Braf*^*V600E*^-positive melanocytes compared to wild-type (Figure 4B).

**Figure 4:**
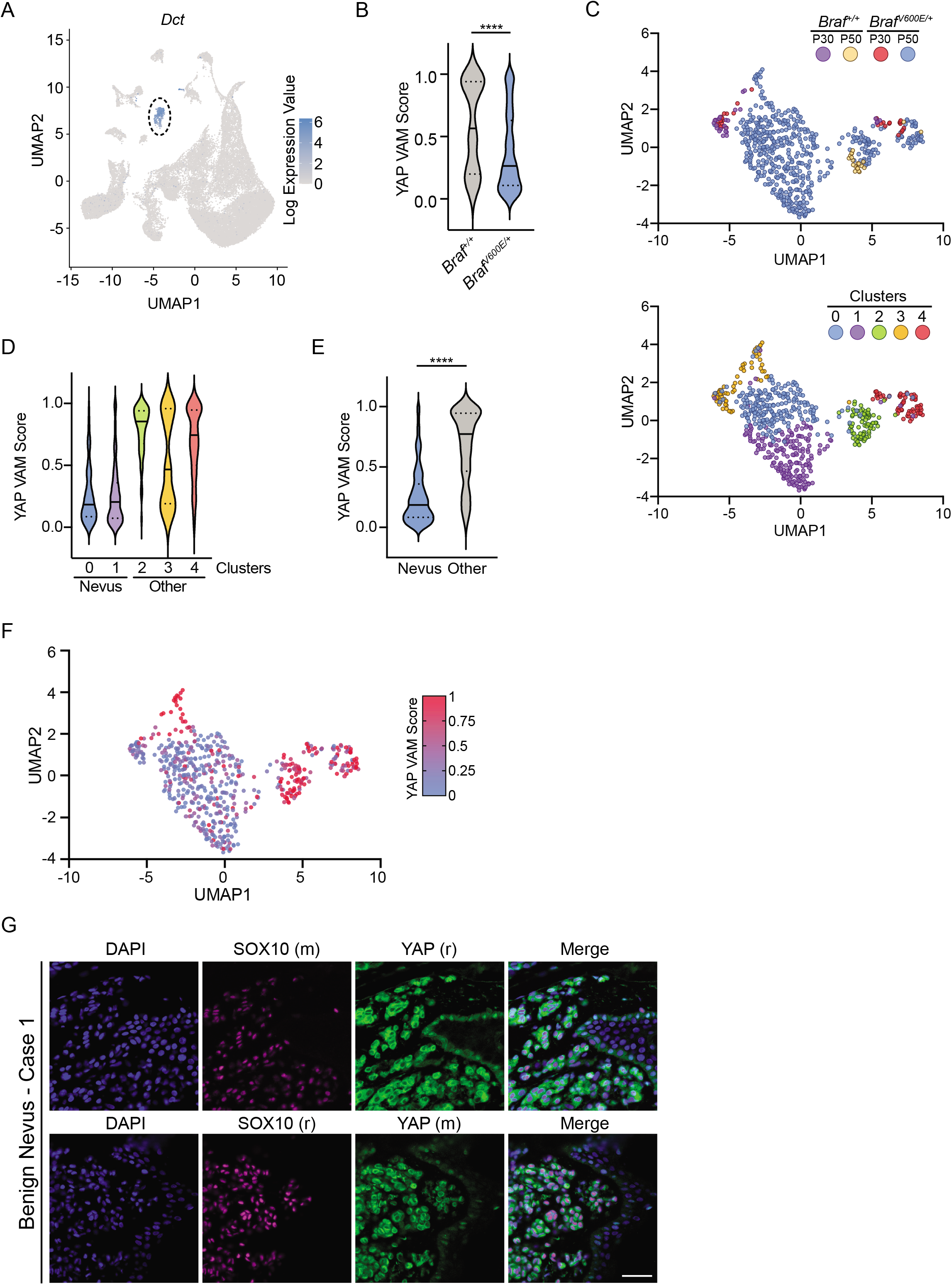
*Braf*^*V600E*^-positive Nevus Melanocytes Display Decreased YAP/TAZ Signaling. **(A)** UMAP of all single-cells from GSE154679 displaying relative expression of melanocyte marker *Dct*. **(B)** Collective YAP VAM score for melanocytes from indicated genotypes (n = 46 for *Braf*^*+/+*^, n = 543 for *Braf*^*V600E/+*^, two-tailed Mann-Whitney test). **(C)** Top, UMAP of melanocytes colored by genotype and animal age; Bottom, UMAP of melanocytes colored by subcluster (n = 589). **(D)** YAP VAM score plotted by melanocyte subcluster. **(E)** YAP VAM score comparing nevus (clusters 0, 1) and other melanocytes (clusters 2, 3, 4) (nevus n = 408, other n = 181, two-tailed Mann-Whitney test). **(F)** UMAP of melanocytes colored by indicated gradient dependent upon YAP VAM score. **(G)** Representative immunofluorescence staining of indicated proteins in a benign nevus with two different sets of antibodies, (r) = rabbit, (m) = mouse, DAPI (blue), YAP (green), SOX10 (magenta), scale bar 50 µm. \*\*\*\**P* < 0.0001.

We then performed unsupervised clustering, which generated five melanocyte subdivisions, to clarify which unique populations of oncogenic *Braf*^*V600E*^*-*expressing melanocytes exhibited the least YAP/TAZ activity (Figure 4C). We theorized that clusters containing nevus melanocytes would be exclusively populated by cells isolated from *Braf*^*V600E*^ mice, and be the primary melanocytic subtype isolated from 50 day old mice. We identified clusters 0 and 1 as *Braf*^*V600E*^-expressing melanocytes isolated from nevi (Figure S4B). In support of this prediction, expression of *Cdkn2a* was found to be the highest in clusters 0 and 1, although *Cdkn2a* read-depth was limited throughout all clusters (Figure S4C). Compared to all other melanocytes, irrespective of genotype, nevus melanocytes (clusters 0 and 1) exhibited the lowest YAP/TAZ activity scores of any cluster, demonstrating that YAP/TAZ-mediated gene expression is restrained following oncogenic *Braf* expression in mouse nevus melanocytes (Figure 4D-F). Importantly, expression of Hippo pathway components remained unchanged regardless of genotype or cluster, suggesting decreased YAP/TAZ signaling was due to Hippo pathway activation, not increased expression of YAP/TAZ regulators (Figure S4D-F).

Interestingly, not all melanocytes captured from *Braf*^*V600E*^ mice exhibited low YAP/TAZ activity scores. Clusters 2 and 4, which contain an appreciable portion of melanocytes from both *Braf*^*+/+*^ and *Braf*^*V600E*^ mice, demonstrated much higher YAP/TAZ activity relative to nevus melanocytes (Figure S4G). However, within these clusters, YAP/TAZ activity was still decreased in *Braf*^*V600E*^ melanocytes compared to wild-type cells. This suggests cell intrinsic mechanisms (e.g. RhoA inactivation) following *Braf*^*V600E*^ expression are only partially leading to decreased YAP/TAZ activity and that other mechanisms, possibly cell extrinsic cues, may play additional roles *in vivo*. We suspect cluster 3, the only cluster that did not exhibit this trend, may be mainly comprised of proliferating, follicular melanocytes as cluster 3 mainly contains melanocytes isolated at P30 when most murine hair follicles are in anagen (25). Taken together, these data reveal *Braf*^*V600E*^*-*expressing mouse melanocytes largely exhibit decreased YAP/TAZ activity, with the most significant decreases found specifically within nevus melanocytes, strongly implying the Hippo pathway becomes activated in response to *Braf*^*V600E*^ expression and nevus formation *in vivo*. In support of these conclusions, immunofluorescence staining of three human benign nevi revealed YAP localization to be predominantly cytoplasmic and thus presumably inactivated in human nevus melanocytes (Figure 4G and S4H).

### *Lats1/2*^*-/-*^ Deletion Promotes Melanomagenesis

Our data suggested that functional inactivation of the Hippo tumor suppressor pathway may enable *Braf*^*V600E*^-expressing melanocytes to evade growth arrest and facilitate melanoma development. To test this, we generated mice carrying floxed alleles of both *Lats1* and *Lats2* (47) with *Tyr::CreER*^*T2*^ to allow for inducible, melanocyte-specific inactivation of the Hippo pathway (*Tyr::CreER*^*T2*^*/Lats1*^*f/f*^*/Lats2*^f/f^). Deletion of *LATS1/2* is well established to completely abrogate the Hippo pathway, and co-heterozygous loss of *LATS1/2* is observed in ∼15% of human melanomas, making deletion of *Lats1/2* clinically relevant (Figure 5A, S5A-B) (2, 48). We also crossed *Tyr::CreER*^*T2*^*/Lats1*^*f/f*^*/Lats2*^*f/f*^ (*Lats1/2*^*-/-*^) mice with mice expressing the Cre-activatable oncogenic *Braf* allele (*Braf*^*CA/+*^), generating *Tyr::CreER*^*T2*^*/Braf*^*CA*^*/Lats1*^*f/f*^*/Lats2*^f/f^ (*Braf*^*V600E*^*/Lats1/2*^*-/-*^) mice (Figure 5B). We confirmed the melanocytic specificity of our *Tyr::CreER*^*T2*^ expressing mice via incorporation of a fluorescent lineage trace (*YFP*^*LSL*^), whose expression was only observed in cells that co-stained for melanocyte markers (Figure S5C).

**Figure 5:**
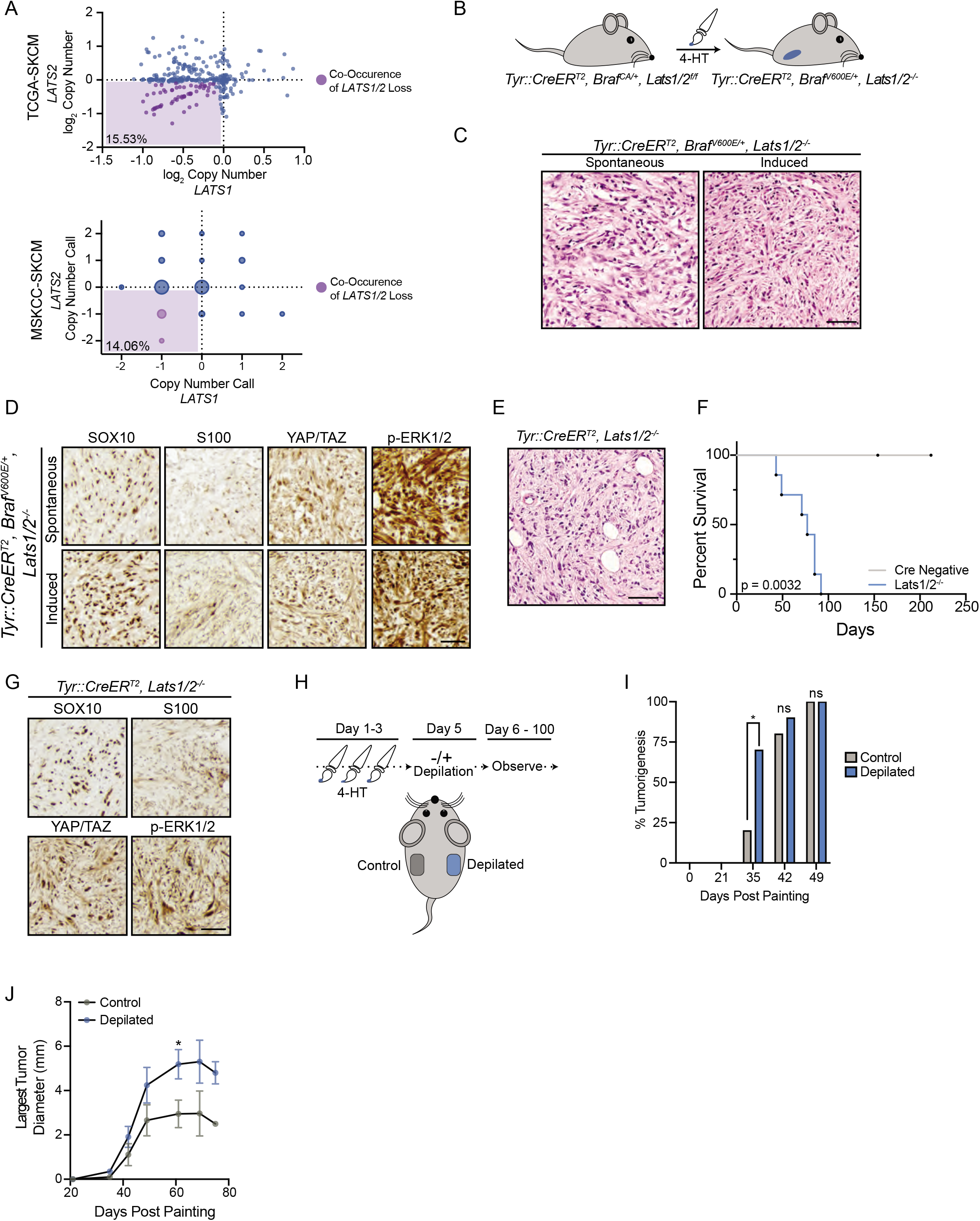
*Lats1/2* Deletion Promotes Melanoma Formation. **(A)** Plot of log_2_ copy number values from TCGA-SKCM or GISTIC 2.0 calls from MSKCC-SKCM databases for the genes *LATS1* and *LATS2*; bottom left percent is frequency of *LATS1/2* co-heterozygous loss. **(B)** Cartoon depicting 4-HT painting experiment. **(C)** Representative hematoxylin and eosin staining of spontaneous or induced tumors from *Braf*^*V600E*^*/Lats1/2*^*-/-*^ mice, scale bar 50 µm (n > 3 spontaneous tumors, n = 2 induced tumors from *Braf*^*V600E*^*/Lats1/2*^*-/-*^ mice). **(D)** Representative IHC of indicated antigens, scale bar 40 µm (n > 3 spontaneous tumors, n = 2 induced tumors from independent *Braf*^*V600E*^*/Lats1/2*^*-/-*^ mice with similar results) **(E)** Representative hematoxylin and eosin staining of induced flank tumor from *Lats1/2*^*-/-*^ mouse, scale bar 50 µm (n > 6 independent mice with similar results). **(F)** Time from painting to a study endpoint of *Lats1/2-/-* mice with 4-HT painted on flanks (Cre negative n = 4, *Lats1/2*^*-/-*^ n = 7, log-rank test). **(G)** Representative IHC of indicated antigens, scale bar 40 µm (n ≥ 3 independent *Lats1/2*^*-/-*^ mice with similar results). **(H)** Cartoon depicting depilation experiment in *Lats1/2*^*-/-*^ mice. **(I)** Percent palpable tumorigenesis in *Lats1/2*^*-/-*^ mice at indicated time points (n = 10 mice, two-tailed unpaired t test). **(J)** Diameter of largest tumor on each animal measured at indicated time points (n ≥ 2 tumors measured at each point, 10 mice total, two-tailed unpaired t test). \**P* < 0.05, ns = non-significant

We observed that *Braf*^*V600E*^*/Lats1/2*^*-/-*^ mice were highly prone to developing spontaneous dermal tumors within weeks after birth, even without topical 4-hydroxytamoxifen (4-HT) administration. A similar melanoma mouse model, *Tyr::CreER*^*T2*^*/Braf*^*CA*^*/Pten*^*f/f*^ (*Braf*^*V600E*^*/Pten*^*-/-*^), is prone to spontaneous melanoma formation in the absence of topical 4-HT, due to leakiness of the inducible Cre recombinase (49, 50). This suggested that deletion of *Lats1/2* was also playing a major role in promoting melanoma development, as *Braf*^*V600E*^ expression alone in murine melanocytes does not generate tumors (5). In the few mice where spontaneous tumorigenesis was absent or delayed, 4-HT administration to *Braf*^*V600E*^*/Lats1/2*^*-/-*^ flanks resulted in the potent formation of tumors which appeared similar to the spontaneously arising neoplasms (Figure 5C). These tumors exhibited strong nuclear YAP/TAZ staining, indicating Hippo inactivation, and positively stained for the melanocytic markers SOX10 and S100 (Figure 5D). SOX10 staining was nuclear and homogenous whereas S100 staining was weakly heterogeneous. Subsequent histopathologic analysis by a board certified dermatopathologist confirmed these infiltrative, spindle cell tumors to be mouse melanoma. Unlike other *Braf*^*V600E*^-driven mouse melanoma models (e.g. *Braf*^*V600E*^*/Cdkn2a*^*-/-*^, *Braf*^*V600E*^*/Trp53*^*-/-*^), which still mainly induce nevus formation, we were unable to appreciate any obvious nevogenesis in *Braf*^*V600E*^*/Lats1/2*^*-/-*^ mice. These data imply oncogenic *Braf*^*V600E*^-positive melanocytes may be incapable of entering an enduring growth arrest without a functional Hippo tumor suppressor pathway.

We also investigated the consequences of *Lats1/2* loss in melanocytes in the absence of oncogenic *Braf*. Intriguingly, we found that following melanocyte-specific deletion of *Lats1/2*, mice exhibited no obvious hyperpigmentation, yet still rapidly developed cutaneous tumors with 100% penetrance after 4-5 weeks (Figure 5E-F, S5D). Co-heterozygous deletion of *Lats1/2* also promoted cutaneous tumorigenesis, albeit at prolonged time scales (Figure S5E). Analysis of *Lats1/2*^*-/-*^ tumor sections revealed non-pigmented neoplasms which were remarkably similar to invasive *Braf*^*V600E*^*/Lats1/2*^*-/-*^ mouse tumors, exhibited a comparable staining profile, and were subsequently diagnosed as mouse melanoma (Figure 5G). Previous studies have identified TEAD and AP-1 transcription factors as major regulators of the melanoma invasive state, which is marked by dedifferentiation and loss of pigmentation signatures (51, 52). We hypothesized that YAP/TAZ-TEAD activation, driven by *Lats1/2* deletion, may enable melanocytes to directly access this invasive gene program explaining our observed lack of pigmentation. In support of this hypothesis, both *Braf*^*V600E*^*/Lats1/2*^*-/-*^ and *Lats1/2*^*-/-*^ mouse melanomas exhibited markedly low staining for mature, differentiated melanocyte markers (Figure S5F).

Given *Lats1/2*^*-/-*^ tumors did not exhibit overt signs of pigmentation, we sought to generate additional data to validate that these neoplasms were melanocytic in origin. It has recently been demonstrated that initiation of the hair follicle cycle, via depilation, strongly promotes melanocyte transformation in *Braf*^*V600E*^*/Pten*^*-/-*^ mice (53, 54). We leveraged this unique characteristic of mouse melanomagenesis to test if *Lats1/2*^*-/-*^ tumor formation was also promoted by depilation, suggesting a melanocytic origin. We induced loss of *Lats1/2* on opposing mouse flanks and then depilated only one flank so as to compare the tumorigenic rate from depilated and non-depilated regions (Figure 5H). We observed that skin regions depilated following 4-HT treatment demonstrated significantly faster tumorigenesis, with palpable tumors observed in 70% of depilated areas compared to 20% of non-depilated areas about one month following treatment (Figure 5I). Not only did tumors appear faster in depilated areas, but these tumors also grew significantly larger (Figure 5J). Collectively, our data demonstrate that melanocyte-specific loss of *Lats1/2* alone, or in conjunction with oncogenic *Braf* expression, promotes melanocyte transformation and melanoma formation *in vivo*.

Next, we investigated whether Hippo pathway inactivation was also occurring in other mouse models of melanoma. We surveyed publicly available datasets (7) and performed gene set enrichment analysis (GSEA) on microarrays gathered in an elegant series of experiments which isolated melanocytes that had either arrested or fully transformed into melanoma in either *Braf*^*V600E*^*/Cdkn2a*^*-/-*^ or *Braf*^*V600E*^*/Cdkn2a*^*-/-*^*/Lkb1*^*-/-*^ mice. GSEA revealed that YAP/TAZ gene sets were significantly enriched in mouse melanoma as compared to arrested, nevus melanocytes or proliferating, non-tumorigenic melanocytes. These data thus indicate that as mouse melanocytes transform into melanoma they exhibit inactivation of the Hippo tumor suppressor pathway (Figures 6A and S6A-D).

**Figure 6:**
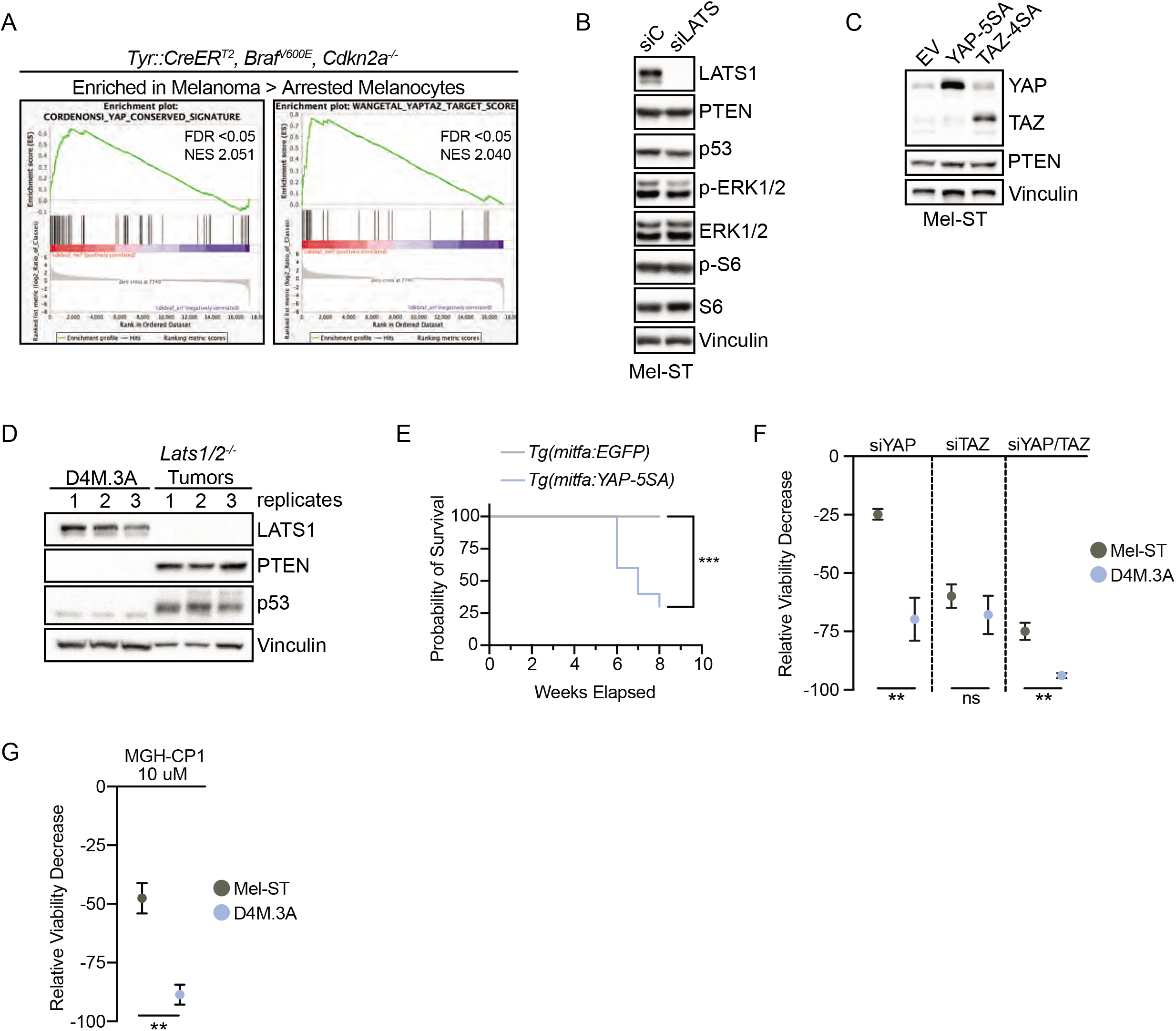
Hippo Pathway Inactivation Promotes Melanoma Formation Across Multiple Models. **(A)** GSEA performed on GSE61750 comparing enrichment of YAP/TAZ signatures in melanoma to arrested melanocytes in indicated genotype (see Figure S6A and S6D). **(B)** Representative IB of Mel-ST parental cell line treated with indicated siRNA (n = 3 independent experiments with similar results). **(C)** Representative IB of Mel-ST parental cell line expressing indicated plasmids, EV: empty vector (n = 3 independent experiments with similar results). **(D)** IB comparing D4M.3A cells and *Lats1/2-/-* tumor lysates where numbers represent replicates or different tumors, respectively (n = 3 independent experiments or mice). **(E)** Survival curve of zebrafish with indicated genotypes (n = 14 for EGFP fish, n = 10 for YAP-5SA fish, log-rank test). **(F)** Percent relative viability decrease in indicated cell lines treated with indicated siRNA for 4 days (n = 3 independent experiments in ≥ technical quintuplicate, graph shows mean ± SEM, two-tailed unpaired t test). **(G)** Relative viability of D4M.3A or Mel-ST cells treated with indicated doses of the TEAD inhibitor MGH-CP1 (n = 3 independent experiments in technical quintuplicate, graph shows mean ± SEM, two-tailed unpaired t test). \*\**P* < 0.01, \*\*\**P* < 0.001, ns = non-significant

### Melanoma Development Can Be Driven by YAP Activation

We sought to further define how deletion of *Lats1/2* promotes melanoma development *in vivo*. While it is well described that *Lats1/2* loss functionally inactivates the Hippo pathway and leads to the activation of YAP and TAZ, LATS1/2 can impinge upon additional signaling pathways that promote tumor development. For example, recent research has revealed inactivation of the Hippo pathway can promote mTOR signaling via multiple routes including YAP-driven expression of a micro-RNA, miR-29, which targets PTEN mRNA for silencing (55-57). However, we detected no observable changes in PTEN protein level following either RNAi-mediated knockdown of LATS1/2 or expression of constitutively active YAP (YAP-5SA) or TAZ (TAZ-4SA) in Mel-ST cells (Figure 6B and C). We could also not appreciate any significant change in phosphorylated S6 levels following *LATS1/2* silencing. Furthermore, examination of *Lats1/2*^*-/-*^ tumors revealed PTEN remained strongly expressed *in vivo* (Figure 6D and S6E). These data reveal that loss of *Lats1/2* is not driving melanomagenesis by activating mTOR via miRNA-mediated depletion of PTEN.

It has also been demonstrated that active LATS2 can bind and inhibit MDM2 leading to increased p53 protein levels (58), raising the possibility that deletion of *Lats1/2* leads to decreases in p53, which may facilitate *Braf*^*V600E*^-driven murine melanomagenesis (59). Discounting this, we found that p53 still accumulates in LATS1/2 depleted Mel-ST cells and *Lats1/2*^*-/-*^ tumors (Figures 6B and 6D).

To assess whether YAP activation alone is sufficient to broadly promote melanoma development, we generated a transgenic zebrafish model that expresses constitutively active YAP specifically in zebrafish melanocytes utilizing the miniCoopR system (60). While all *EGFP*-expressing controls experienced no tumor burden in this system, all *Tg(mitfa:YAP-5SA*) zebrafish rapidly developed pigmented fish melanoma (Figure 6E). This result demonstrates that constitutively active YAP is sufficient to induce melanoma development, and highlights *YAP1* as an important melanoma oncogene. Although deletion of *Lats1/2* may still be promoting mTORC1 activity in our murine model, given activation of mTORC1 alone is insufficient to promote melanoma formation *in vivo* (7), whereas activation of YAP alone is sufficient, we believe activation of YAP/TAZ remains essential for the observed tumor formation in both *Braf*^*V600E*^*/Lats1/2*^*-/-*^ and *Lats1/2*^*-/-*^ mice.

As activation of YAP/TAZ signaling was observed to promote tumorigenesis across *in vivo* melanoma models, we investigated whether depletion of YAP/TAZ could inhibit melanoma cell growth. RNAi-mediated knockdown of YAP/TAZ in a *Braf*^*V600E*^*/Pten*^*-/-*^ mouse melanoma cell line (D4M.3A) resulted in significantly decreased viability as compared to immortalized melanocytes (Figure 6F). This decrease in viability was predominantly driven by loss of YAP, as single knockdown of YAP was significantly detrimental to *Braf*^*V600E*^*/Pten*^*-/-*^ melanoma cells (Figure 6F). To further confirm loss of YAP/TAZ signaling was driving decreases in viability, we treated either Mel-ST or D4M.3A cells with a recently discovered pan-TEAD inhibitor, MGH-CP1 (61). Strikingly, *Braf*^*V600E*^*/Pten*^*-/-*^ mouse melanoma cells were exquisitely sensitive to inhibition of TEAD by MGH-CP1 (Figure 6G). Moreover, combined inhibition of TEAD and MAPK signaling further decreased D4M.3A cell viability (Figure S6F).

Together, these data implicate YAP as a cutaneous melanoma oncogene and novel therapeutic target to further explore in the treatment of human melanoma. While *YAP1* is not commonly mutated in human melanoma, *YAP1* amplifications and mutations have been observed, and YAP staining in primary melanoma has been shown to significantly correlate with reduced patient survival (31, 62, 63).

## Discussion

Discerning the molecular pathways that govern the growth arrest of nevus melanocytes, and how melanocytes ultimately overcome these barriers, is critical to fully understanding the mechanisms of melanomagenesis. A considerable body of work supports a role for OIS in preventing tumorigenic growth of melanocytes *in vitro* and *in vivo*; however, expanding lines of evidence demonstrate melanocytes within nevi retain proliferative capacity, in conflict with the absolute growth arrest implied by OIS (16). Indeed, up to 30% of melanomas are predicted to arise from pre-cursor nevi (17). These observations highlight nevi as being the product of “stable clonal expansion”, rather than senescence (16).

Several genetic alterations that have the capacity to overcome the stable growth arrest of melanocytes expressing oncogenic MAPK mutations have been identified, including *CDKN2A, PTEN*, and *TP53* (3, 5-7, 59). While deletion of *Cdkn2a* and *Trp53* with *Braf*^*V600E*^ expression induces murine melanomagenesis, the vast majority of melanocytes still enter a stable growth arrest to form nevi (7). Further, recent murine single-cell sequencing analyses reveal *Braf*^*V600E*^-positive nevus melanocytes do not exhibit expression of senescence signatures (25). Together, these discoveries suggest additional unidentified tumor suppressive mechanisms exist to enforce the arrest of nevus melanocytes.

Our data demonstrate that the Hippo pathway represents one such tumor suppressive mechanism. We discovered that expression of *BRAF*^*V600E*^ promotes activation of the Hippo tumor suppressor pathway across multiple cell lines and that *Braf*^*V600E*^-positive murine nevus melanocytes display significantly decreased YAP/TAZ signaling *in vivo*. We determined oncogenic *BRAF* expression induces Hippo pathway activation and cell cycle arrest *in vitro* via a cell-intrinsic mechanism, in which hyperactive MAPK signaling alters the cytoskeleton in part through decreased RhoA signaling. By contrast, our analysis of single-cell sequencing data indicated that while *Braf*^*V600E*^-expression leads to a general repression of YAP/TAZ signaling in most melanocytes *in vivo*, the effect is strongest in nevus melanocytes (Figure 4D). This suggests cell-extrinsic cues, perhaps specific to the nevus microenvironment, may also be impinging upon YAP/TAZ signaling.

A recent modeling study has proposed that nevus melanocyte growth arrest is likely governed by collective cell behaviors (e.g. paracrine signaling (10)), rather than cell-autonomous mechanisms (25). Highly analogous to the collective cell processes mediating contact inhibition and organ growth, which are governed by the Hippo pathway, it is well known that a majority of *Braf*^*V600E*^-expressing melanocytes only enter a state of growth arrest following multiple rounds of division. It is therefore tempting to speculate that melanocyte overgrowth induced by oncogene expression significantly increases melanocyte density over multiple rounds of division ultimately promoting mounting activation of the Hippo tumor suppressor pathway and eventual growth arrest. If true, one would predict Hippo pathway activation would only engage after a nevus passes a critical size threshold and that nevus melanocyte growth may be restored by reducing local melanocyte density. Intriguingly, nevi that are partially resected have been observed to regain proliferative capacity and the vast majority of these recurrent nevi did not grow beyond the limits of the original surgical scar, suggesting these nevus melanocytes arrest once they attain a similar size (64, 65).

We also found that Hippo pathway impairment alone is sufficient to induce melanoma formation in various *in vivo* models, even upon heterozygous loss of *Lats1/2*. This finding is clinically relevant, as co-heterozygous loss of *LATS1/2* is frequently observed in human melanomas (Figure 5A). However, it should be noted that Hippo pathway inactivation can also occur through mechanisms independent of *LATS1/2* loss. For example, >80% of uveal melanomas and ∼6% of cutaneous melanomas have mutations in *GNAQ/GNA11* which can activate YAP/TAZ independent of LATS1/2 *(66-68)*. Intriguingly, these activating mutations in *GNAQ*/*GNA11* have been shown to stimulate RhoA activity (66). As YAP/TAZ-TEAD activation appears to be relatively common in melanoma, it will be important to define additional mechanisms by which Hippo inactivation occurs (63, 69).

Our live-cell imaging revealed induction of oncogenic *BRAF* promotes chromosome alignment defects, prolonged mitosis, and a significant increase in cells experiencing a whole-genome doubling (WGD) event. We predominantly detected mitotic slippage events, in which cells expressing *BRAF*^*V600E*^ exited mitosis without undergoing cell division, thus giving rise to multinucleated tetraploid cells which are commonly observed within benign human nevi (37). Typically caused by mitotic or cytokinetic failures, cells that undergo a WGD event (referred to as WGD^+^) are genomically unstable cells known to drive tumorigenesis, and their contribution to human cancer is significant (70-72). In fact, the vast majority of metastatic melanomas and ∼30-40% of human melanomas overall show evidence of having experienced a WGD event (71-73). WGD^+^ cells are known to activate both the Hippo and p53 tumor suppressor pathways to restrain their ongoing proliferation (35, 58, 74, 75). As such, Hippo pathway inactivation would restore proliferation not only to growth-arrested, diploid, nevus melanocytes, but also to the rarer, possibly WGD^+^, multinucleated melanocytes. Whether the ∼40% of melanomas that have experienced a WGD enrich for alterations in the Hippo pathway and what specific defects cooperate with oncogenic MAPK mutations to promote WGD in human melanoma remains unknown.

Questions remain as to how Hippo pathway activation so strongly promotes melanoma development. For example, activating MAPK mutations appear to be critical for human melanocyte transformation. However, we establish that mouse melanocytes merely lacking *Lats1/2* rapidly develop melanoma. Interestingly, while *Lats1/2* loss does not stimulate the MAPK pathway *in vitro*, we found that *Lats1/2*^-/-^ melanomas exhibit strong p-ERK staining (Figure 5G). These data suggest that *Lats1/2*^-/-^ melanocytes hyperactivate the MAPK pathway either through the acquisition of genetic alterations or, potentially, deletion of *Lats1/2 in vivo* is capable of stimulating MAPK activity through YAP/TAZ-dependent transcriptomic changes (67). Indeed, it has been observed that one common resistance mechanism to Vemurafenib, a BRAF^V600E^-specific inhibitor, is the amplification and/or activation of YAP (42, 69, 76). This suggests that YAP activity can compensate for loss of *BRAF*^*V600E*^ signaling. Given MAPK inhibitor resistance remains a significant component of treatment failure, our data suggests that co-targeting MAPK and YAP-TEAD signaling could simultaneously prevent resistance (69) and decrease melanoma cell viability (32, 77).

In summary, we demonstrate that activation of the Hippo tumor suppressor pathway promotes melanocyte growth arrest in response to the expression of oncogenic *BRAF*^*V600E*^. Further, disruption of the Hippo pathway, which promotes activation of YAP/TAZ, potently induces melanocyte growth *in vitro* and melanoma development *in vivo* in multiple model organisms. Collectively, our data implicate the Hippo pathway as an important melanoma tumor suppressor and highlight YAP/TAZ as promising therapeutic targets to investigate for the treatment of human melanoma.

## Acknowledgments

We would like to thank Jackie King and Ross King for their unwavering resolve to promote melanoma awareness and research, as well Qi Sun, Shuyang Chen, and the entire Ganem Lab for wisdom and advice. We would also like to thank April Deng, Karen Dresser, Constance Brinckerhoff, Robert Weinberg, Craig Ceol, and Arthur Lander for sharing cell lines, reagents, and/or assistance. The results published herein are partly based upon data generated by the TCGA Research Network: https://www.cancer.gov/tcga. We thank all the patients who donated specimens to both the TCGA and the MSKCC database. M.A.V. is supported by a Ruth L. Kirchstein National Research Service Award (F30) from the National Cancer Institute (1F30CA228388). E.X. and S.M. were both supported by an award from the Boston University Undergraduate Research Opportunities Program. X.V. is supported by NIH NHLBI (R01HL124392) and an American Cancer Society -Ellison New England Research Scholar Grant (RSG-17-138-01-CSM). N.M.K was supported by NIH NHLBI grant F31HL146163. N.J.G. is a member of the Shamim and Ashraf Dahod Breast Cancer Research Laboratories and is currently supported by NIGMS (GM117150), and the Harry J. Lloyd Charitable Trust. N.J.G. was previously supported by the Jackie King Young Investigator Award from the Melanoma Research Alliance and the Searle’s Scholars Program.

## Author Contributions

M.A.V. and N.J.G conceptualized the study, designed the *in vitro* experiments, and wrote the manuscript. M.A.V., N.K, X.V, and N.J.G designed all *in vivo* studies. M.A.V. performed most of the cell biological assays, tissue staining, and imaging analysis. N.K. performed *in vivo* experiments and prepared all tissue samples. N.K., E.X., S.M., and A.T.L. assisted M.A.V. with the cell biological and tissue assays. X.X. provided dermatopathology consult and tissue analysis. R.D. and C.C. performed zebrafish experiments. R.H. and J.C. completed the single-cell analysis and M.A.V. performed GSEA. L.H. and D.L. provided critical reagents.

## Materials and Methods

### Cell Culture

Immortalized melanocyte (Mel-ST) cells were a gift from the lab of Dr. Robert Weinberg. Mel-ST cells, and all derivative cell lines generated in this study, were grown in DMEM media containing 5% Fetal Bovine Serum (FBS), 100 IU/mL penicillin, and 100 µg/mL streptomycin. Human Embryonic Kidney 293 (HEK293A) cells, and all derivative cell lines, were grown in DMEM media containing 10% FBS, 100 IU/mL penicillin, and 100 µg/mL streptomycin. hTERT-BJ fibroblasts, and all derivative cell lines, were grown in DMEM:F12 media containing 10% FBS, 100 IU/mL penicillin, and 100 µg/mL streptomycin. *Braf*^*V600E*^*/Pten*^*-/-*^ mouse tumor cells (D4M.3A) were a generous gift of Dr. Constance Brinckerhoff. D4M.3A mouse tumor cells were maintained in DMEM:F12 media containing 5% FBS. Primary adult epidermal melanocytes were purchased from ATCC (PCS-200-013) and maintained in Dermal Cell Basal Medium (ATCC PCS-200-030) supplemented with an Adult Melanocyte Growth Kit (ATCC PCS-200-042). All FBS used in these studies was confirmed to either be naturally absent of tetracyclines or below 20 ng/mL by the manufacturer. All cells were maintained at 37°C with 5% CO_2_ atmosphere and maintained at sub-confluent levels for passaging and all experiments. Cultures were regularly checked for mycoplasma contamination utilizing a PCR detection kit (G238, ABM) or Hoechst staining. Bright-field images of tissue culture cells were captured on an Echo Revolve Hybrid Microscopy system at 10X or 20X (Echo Laboratories).

### Cell Line Generation

To generate the *BRAF*^*V600E*^ doxycycline inducible system Mel-ST, HEK293A, or hTERT BJ fibroblasts were first infected with lentivirus generated from pLenti CMV TetR Blast (Tet Repressor) and selected. Following selection, cells were infected with lentivirus generated from pLenti CMV/TO BRAF^V600E^ Neo or pLenti CMV/TO BRAF Neo, selected, and single cell cloned to establish cell lines which demonstrated no basal expression at baseline and strong induction after doxycycline addition. Expression of *BRAF*^*V600E*^ was confirmed using two different mutant-specific antibodies (VE1 and RM8 clones). To generate stably expressing H2B-GFP lines, cells were infected with lentivirus generated from pLenti H2B-GFP Blast. Stably expressing Mel-ST cells with empty vector (pLVX Puro), YAP-5SA (pBABE YAP-5SA Puro), and TAZ-4SA (pLVX Flag TAZ-4SA Puro) were generated via viral infection followed by selection. Tetraploid cells were generated by treating asynchronous cells with 4uM DCB for 16 h, followed by gentle washing to remove drug (5 × 5 min); completion of cytokinesis was confirmed by phase-contrast imaging.

### Viral Infections and Transfections

Mel-ST, HEK293A, or BJ Fibroblasts were infected for 12-16 h with virus carrying genes of interest in the presence of 10 µg/mL polybrene, washed, and allowed to recover for 24 h before selection or single cell cloning. Short term viral infection of primary melanocytes was carried out similarly, but with 2 µg/mL of polybrene. All RNAi transfections were performed using 25-50 nM siRNA with Lipofectamine RNAi MAX according to the manufacturer’s instructions. Briefly, cells were seeded in 6 or 12-well plates either by addition of reverse transfection mixture overnight, which was then washed and replaced with fresh media 18 h later, or forward transfection mixture for 4 h, which was then replaced with fresh media. Cells were then incubated for 48-72 h prior to lysis at sub-confluent levels.

### Plasmid Generation

Plasmids encoding the Tetracycline Repressor, pLenti CMV TetR Blast (716-1), was a gift from Eric Campeau & Paul Kaufman (Addgene Plasmid #17492). To create pLenti CMV/TO BRAF Neo and pLenti CMV/TO BRAF^V600E^ Neo, we performed Gateway cloning using Gateway LR Clonase II (Invitrogen) according to manufacturer instructions to insert BRAF or BRAF^V600E^ into pLenti CMV/TO Neo DEST (685-3) using pENTR BRAF or pENTR BRAF^V600E^. pENTR BRAF and pENTR BRAF^V600E^ were gifts from Craig Ceol and pLenti CMV/TO Neo DEST (685-3) was a gift from Eric Campeau & Paul Kaufman (Addgene plasmid #17292). To generate pLenti CMV/TO NRAS^Q61R^ Neo we performed Gateway cloning to insert NRAS^Q61R^ from the donor vector pDONR223 NRAS^Q61R^ into the destination vector pLenti CMV/TO Neo Dest. pDONR223 NRAS^Q61R^ was a gift from Jesse Boehm, William Hahn, and David Root (Addgene plasmid #81652).

### Immunofluorescence and Confocal Microscopy

Cells were plated on glass coverslips, treated as indicated, washed in 1X phosphate-buffered saline (PBS) (Boston Bioproducts) and fixed in 4% paraformaldehyde for 10 min. Cells were then washed in PBS-0.01% Triton X-100, extracted in PBS-0.2% Triton X-100 for 10 min, blocked in Tris-buffered saline (TBS)-bovine serum albumin (BSA) (10 mM Tris, pH 7.5, 150 mM NaCl, 5% Bovine Serum Antigen, 0.2% sodium azide) for 1h, and incubated with primary antibodies diluted in TBS-BSA for 1 h at RT or overnight at 4°C in a humidified chamber. Primary antibodies were visualized using species-specific fluorescent secondary antibodies (Molecular Probes, Alexa Fluor secondaries, 488 nm, 568 nm, 1:500) and DNA was detected with 2.5 µg/mL Hoechst. F-actin was visualized using rhodamine-conjugated phalloidin (1:2000, Molecular Probes, R415). Immunofluorescence images for analysis were collected on a Nikon Ti-E inverted microscope equipped with a Zyla 4.2 PLUS (Andor) and X-Cite 120 LED light source at the same exposure. Confocal immunofluorescence images were collected on a Nikon Ti-E inverted microscope equipped with a C2+ laser scanning confocal head with 405 nm, 488 nm, 561 nm, 640 nm laser lines. Z-stacks were acquired with a series of 0.5-1 µm optical slices which were then converted into a single, max-intensity projected image. Images were analyzed using NIS-Elements Advanced Research (AR) and ImageJ. To assess YAP localization, two small square regions of interest were drawn at random in individual cells with one in the nucleus, and one in the cytoplasm. The background corrected, mean fluorescence intensity of YAP was subsequently measured in these regions of interest and a nuclear to cytoplasmic ratio was determined. To assess stress fiber quantity, images were background corrected, contrast normalized, and then fibers obvious to the naked eye were counted.

### Live-Cell Imaging

Stably expressing H2B-GFP cells were grown on glass-bottom 12-well tissue culture treated dishes (Cellvis) and treated with drugs of interest. Immediately post-treatment imaging was performed on a Nikon Ti-E inverted microscope equipped with the Nikon Perfect Focus system. The microscope stage was enclosed within a temperature and atmosphere-controlled environment at 37°C and 5% humidified CO_2_. Fluorescent or bright-field images were captured every 5-10 min with an 10X or 20X 0.5 NA Plan Fluor objective at multiple locations for 72-96 h. All captured images were analyzed using NIS-Elements AR software. Mitotic length was calculated by counting the duration from nuclear envelope breakdown to anaphase onset.

### Tissue Staining

At the experimental endpoint mice were euthanized and mouse tumors or skin samples were dissected and immediately fixed in 4% paraformaldehyde (PFA) in PBS for 16h at 4°C. Tissues were then paraffin embedded for sectioning and mounting. PFA-fixed paraffin-embedded tissue sections were cut at 5 µm and mounted onto positively charged coverslips (Colorfrost Plus, Thermo). Mounted tissue samples were deparaffinized using xylenes and rehydrated via an ethanol:water gradient. For hematoxylin and eosin staining (H&E), the tissues were incubated in hematoxylin (#14166, Cell Signaling Technology), rinsed in water, differentiated with 1% acid ethanol, blued with 0.1% sodium bicarbonate solution, rinsed in water, dehydrated, cleared, and mounted with Cytoseal XYL (Thermo Scientific). For immunohistochemistry (IHC) or immunofluorescence (IF), antigen unmasking was performed on rehydrated tissue sections using either a citric-acid retrieval buffer (Vector Labs) or Tris-EDTA retrieval buffer (10 mM Tris, 1 mM EDTA, 0.05% Tween-20). Heat-mediated antigen retrieval was performed using either a standard microwave (95°C, 20 min) or Decloaking Chamber NxGen (Biocare Medical) (110°C, 12 min), followed by cooling to room temperature (∼30-60 min). Citric acid retrieval was used for most antibodies; Tris-EDTA retrieval was used for MelanA antibodies. For IHC, tissue sections were washed, endogenous peroxidase activity was quenched using 3% hydrogen peroxide in PBS for 10 min, and then blocked for 1h in 10% goat serum in TBS (Sigma-Aldrich). For IF, tissue sections were blocked for 1h in 10% goat serum in TBS. Following serum block, if necessary, tissue was incubated with Rodent Block M (Biocare Medical) for 30 min to block endogenous mouse IgG prior to murine primary antibody addition. Primary antibodies were diluted in 10% goat serum in TBS and incubated overnight at 4°C in a humidified chamber. Following primary addition for IHC, slides were washed with TBS-0.01% Tween-20, incubated with anti-rabbit or mouse SignalStain Boost IHC detection reagent (Cell Signaling Technology) for 30 min and then developed with SignalStain DAB substrate kit (Cell Signaling Technology) according to manufacturer’s instructions. Counterstaining was performed using hematoxylin, followed by dehydration, clearing, and mounting with Cytoseal XYL (Thermo Fisher). Images were captured at randomly selected points using a Nikon Ti-E inverted microscope equipped with a DS-Ri2 (Nikon). For IF, slides were incubated with species-specific fluorescent secondary antibodies (Molecular Probes) and 2.5 µg/mL Hoechst for 1h at room temperature in a dark humidified chamber. Auto-fluorescence was quenched using Vector TrueVIEW according to manufacturer’s instructions, and slides were mounted using Prolong Gold Antifade (Invitrogen). Images were captured at randomly selected points using an Nikon Ti-E inverted microscope equipped with a Zyla 4.2 PLUS (Andor) and X-Cite 120 LED light source. For all staining experiments tissue-specific secondary controls were included in each staining experiment to ensure specificity and control for endogenous tissue pigmentation levels.

### Protein Extraction and Immunoblotting

Cells were gently washed twice in ice-cold 1X PBS and lysed using ice-cold cell lysis buffer (50 mM Tris-HCl, 2% w/v SDS, 10% glycerol) containing 1X HALT (dual phosphatase and protease inhibitor, Thermo Fisher). Lysates were sonicated at 20% amplitude for 20 seconds, diluted in 4X Sample Buffer (Boston BioProducts), and resolved using SDS gel electrophoresis. For mouse tumor samples, 10 mg of tumor tissue was dissected and placed into RIPA lysis buffer (Boston BioProducts) supplemented with one Pierce protease inhibitor tablet and 1X HALT (Thermo Fisher). Tissue was mechanically dissociated initially using a 7cm pestle (Kimble) followed by further homogenization with a 20-guage needle and brief sonication for 20 s at 20 kHz. Tissue lysate was then centrifuged at 4°C and >15,000g for 10 min. Supernatant was collected, diluted in 4X Sample Buffer to 2X, and resolved using SDS gel electrophoresis. For phos-tag immunoblots, a phos-tag gel was prepared with diluted Phos-Tag™ Acrylamide reagent (Wako Chemicals, AAL-107) according to manufacturer’s instructions utilizing manganese as the divalent cation. Proteins were then transferred onto PVDF membranes using a wet-tank transfer system (Bio-Rad), blocked for 1 h with TBS-0.1% Tween-20 containing 5% skim milk powder and then probed overnight at 4°C with primary antibodies diluted in TBS-0.1% Tween-20 containing 1% skim milk powder. Bound antibodies were detected by incubating membranes with horseradish peroxidase-linked, species-specific, secondary antibodies (1:5000, Cell Signaling Technology) for 1 h, followed by 3×10 min washing in TBS-0.1% Tween-20, and then addition of Clarity or Clarity Max ECL blotting substrate (Bio-Rad). Chemiluminescence acquisition was carried out using the Bio-Rad ChemiDox XRS+ system and quantitative densitometry was measured using Image Lab (Bio-Rad).

### Human Nevus Sample Collection

Human skin tissues with melanocytic nevi and melanoma were retrieved from existing material in the pathology laboratory at UMass Medical Center. The tissue blocks are deidentified with new numbers before tissue sections (10 µm) were obtained for IHC stains. These studies were reviewed and determined to be IRB exempt by Boston University School of Medicine.

### RNA isolation and qRT-PCR

Total RNA from cultured cells was isolated using a Quick-RNA kit (Zymo Research). cDNA libraries were generated from RNA using the Superscript III kit and random hexamer primers (Invitrogen). Quantitative real-time PCR was performed using SYBR Green reagents in a StepOnePlus system (Applied Biosystems) according to manufacturer protocol. For each individual experiment a technical triplicate was run which was then averaged to generate a single biological replicate. Primer sequences were as follows: *CYR61*: Forward, AGCCTCGCATCCTATACAACC, Reverse, TTCTTTCACAAGGCGGCACTC *AMOTL2*: Forward, TTGGAATCTGCAAATCGCC, Reverse, TGCTGTTCGTAGCTCTGAG *GAPDH*: Forward, GAGTCAACGGATTTGGTCG, Reverse, CATTGATGGCAACAATATCCAC

### Antibodies

The antibodies used herein categorized by technique and company. Immunofluorescence (cells): Santa Cruz Biotechnology: YAP 63.7, 1:250 (detects both YAP/TAZ) Immunofluorescence (tissue): Cell Signaling Technologies: YAP (1A12) #12395, 1:100; YAP/TAZ (D24E4) #8418, 1:100; YAP (D8H1X) #14074, 1:100; Abcam: gp100 (ab137078), 1:250 Immunohistochemistry: Cell Signaling Technologies: YAP/TAZ (D24E4) #8418, 1:100; YAP (D8H1X) #14074, 1:100; GFP (D5.1, cross reacts with YFP) #2956, 1:100; phospho-p44/42 ERK1/2 (Thr202/Tyr204) #9101, 1:100; Abcam: gp100 (ab137078), 1:250; SOX10 (ab180862), 1:250; MelanA (ab210546), 1:500; Dako: S100 (IS504), pre-diluted by manufacturer. Immunoblotting: Cell Signaling Technologies: B-Raf (D9T6S), 1:1000; phospho-p44/42 ERK1/2 (Thr202/Tyr204) #9101, 1:1000; p-44/42 MAPK (ERK1/2) #9102, 1:1000; RSK1/RSK2/RSK3 (3D27) #9355, 1:1000; phospho-p90RSK (Ser380) (D3H11) #11989, 1:1000; GAPDH (14C10) #2118, 1:1000; YAP (D8H1X) #14074, 1:1000; LATS1 (C66B5) #3477, 1:1000; phospho-LATS1 (Thr1079) (D57D3) #8654, 1:1000; TAZ (E8E9G) #83669, 1:1000; TAZ (V386) #4883, 1:1000; phospho-S6 (Ser235/236) (D57.2.2E) XP #4858, 1:1000; PTEN (138G6) #9559, 1:1000; p53 (1C12) #2524, 1:1000; phospho-Chk1 (Ser345) #2341, 1:500 Abcam: Vinculin (ab18058), 1:4000; Fisher Scientific: BRAF^V600E^ (RM8 Clone) #MA5-24661, 1:1000-2000; Bethyl Laboratories: LATS1/LATS2 (A300-479A), 1:1000 Santa Cruz Biotechnology: Chk1 (G-4) sc-8408, 1:500; S6 (C-8) sc-74459, 1:1000; N-Ras (F155) sc-31, 1:1000 Spring Biosciences: BRAF^V600E^ (VE1 Clone) # E19290, 1:1000

### Soft Agar Assays

In 6-well dishes, sterile 2% noble agar stock solution in water was dissolved by heating to ∼40-45°C, mixed with warm media, plated at final concentration of 0.6%, and allowed to cool and solidify at 4°C. Following solidification gels were warmed back to 37°C. Next, cells were trypsinized, counted, 1×10^4^ cells were plated in 0.3% noble agar and allowed to solidify at room temperature or briefly at 4°C. Plates were maintained in a cell culture incubator for 2-4 weeks with feedings of 1.5 mL of 0.3% agarose solution weekly. All drugs were maintained at 2X concentration in underlays and independent experiments were done in technical triplicate. Total colonies per well were counted using phase-contrast imaging on an Echo Revolve Hybrid Microscopy system at 10X (Echo Laboratories). For imaging, gels were stained for 20 min with 0.1% Crystal Violet, gently washed multiple times overnight, and imaged on a Chemi-Doc XRS+ system under Fast Blast with 0.015-0.1 sec exposure.

### Drug Treatments

The concentrations used for the MAPK pathway inhibitors (MEKi-1/2 and ERKi) were determined experimentally to be the doses at which phosphorylation of ERK1/2 and RSK1/2/3 returned to baseline despite the presence of oncogenic *BRAF* expression. The reagents used in these studies are as follows: MEKi-1: U0126 (Selleck Chemicals), 10 µM; MEKi-2: Trametinib (GSK1120212), 20 nM (Selleck Chemicals); ERKi: SCH772984, 20 nM (Selleck Chemicals); Hydroxyurea, 1 mM (Selleck Chemicals); doxycycline, 1 µg/mL (Sigma-Aldrich D9891); ROCKi: Y-27632, 5-10 µM (Selleck Chemicals); RACi: NSC 23766, 25-50 µM (Selleck Chemicals); thymidine, 2.5 mM (Sigma-Aldrich T1895); RO-3306, 7 µM (Sigma-Aldrich SML0569)

### Population Doubling and Viability Assays

For population doubling assays, initially 1×10^5^ cells (c_initial_) were plated in 10 cm dishes. After 4 days of growth cells were trypsinized, counted, and 1×10^5^ cells were plated again. Cells were trypsinized and counted again after 4 more days. The number of population doublings (pd) were calculated by inputting the counted number of cells (c _final_) into the following equation: 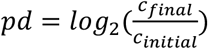. For viability assays, 4×10^3^ cells per well were plated into a white-bottom 96-well dish in technical duplicate to quintuplicate, dependent upon experiment. Cells were then treated with indicated siRNA and/or drugs for 4 days. The 96-well plate was then allowed to equilibrate at room temperature for 30 minutes followed by addition of Cell Titer Glo (Promega) to each well according to manufacturer instructions. Viability was then assessed via luminescence using a BMG microplate reader and analyzed using BMG Labtech Optima v2.0R2 software. All technical replicates were averaged to generate a single biologic replicate.

### Copy Number Analysis

Log_2_ copy-number values and/or putative copy-number GISTIC 2.0 values for *LATS1* and *LATS2* were downloaded from cBioPortal for the TCGA skin cutaneous melanoma (SKCM) subset of the PanCancer Atlas and MSKCC, NEJM 2014 datasets. SKCM abbreviation represents skin cutaneous melanoma. Mutational status for *BRAF* and *NRAS* from the TCGA-SKCM dataset was also obtained from cBioPortal. Graphs were created in Prism 9.

### Gene Expression Analysis

Publicly available microarray dataset GSE61750 was obtained from the Gene Expression Omnibus (GEO). For GSEA indicated groups were normalized via meandiv, probes collapsed to max probe, and a weighted enrichment metric (log_2_ Ratio of Classes) was utilized with 1000 permutations to determine enrichment in indicated gene sets. The YAP/TAZ target score gene set was derived from (45) and (46). Gene set can be found in Supplementary Table 1. The Hippo component gene set was derived from Reactome with additional genes added and can be found in Supplementary Table 2.

### Single-Cell RNAseq Analysis

Count matrix of GSE154679 was downloaded from the GEO repository (25). Cells that had less than 200 expressed genes, more than 4000 expressed genes or > 17% of mitochondrial gene were removed from analysis. A total of 35214 cells were retained for downstream analysis. Normalization, dimensionality reduction, cell-clustering and data visualization were analyzed with Seurat package (78). PCA dimension reduction was performed using top 2000 highest variance genes. The top 15 principal components were utilized to calculate the k-nearest neighbors of each cells. Cell clusters were determined using Louvain algorithm at a resolution = 0.5, which was high enough to obtain clusters associated with cell lineage identity. We used UMAP to visualize and confirm cell clustering. Melanocyte cell population was identified based on the expression of canonical gene markers. To identify subclusters within melanocyte cell populations, we re-analyzed cells within melanocyte cell clusters using the same analysis pipeline described above. To quantify gene sets activity in each cell, gene set testing of scRNA-seq data was performed following VAM pipeline (44). Data Normalization and highest variable genes were analyzed using pipeline described above. VAM method was executed using vamForSeurat() function on melanocyte cell population for each gene set. Graphs were created in Seurat or data was imported into Prism 9 for graph creation.

### Murine Studies

All animal experiments were conducted according to protocols approved by the Institutional Animal Care and Use Committee (IACUC) at Boston University (Protocol # PROTO201800236). All other mouse strains were sourced from Jackson Laboratory: *Tyr::CreER*^*T2*^ (Jax # 012328), *Lats1*^*f/f*^ (Jax # 024941), *Lats2*^*f/f*^ (Jax # 025428), *BRAF*^*V600E*^ (Jax # 017837), *R26-YFP*^*LSL*^ (Jax # 006148). 4-hydroxytamoxifen (4-HT [Sigma, H7904]) was dissolved in methanol to a concentration of 5 mg/ml. To induce knockout in *Lats1/2*^*-/-*^, and *Braf*^*V600E*^*/Lats1/2*^*-/-*^, mice aged 8-12 weeks had an approximately 2 cm square shaved on the right flank using surgical clippers and sufficient 4-HT solution to wet the skin was applied once daily to the shaved area for three consecutive days. For *Braf*^*V600E*^ mice, a higher concentration of 4-HT was required; 4-HT was dissolved in dimethylsulphoxide (DMSO) to a concentration of 25 mg/ml, and topical administration was performed as above. For the depilation experiments using *Lats1/2*^*-/-*^ mice, a 2 cm square was shaved on both the left and right flank of each mouse, and 5 mg/ml 4-HT was topically applied to each square for three consecutive days as above on the right flank, as well as on an approximately 2 cm length of the tail for three consecutive days. Two days after the final application, chemical depilation was performed by application of Nair hair remover onto the right flank for 15 seconds, followed immediately by wiping with a damp tissue to prevent irritation. For systemic knockout experiments, mice were injected with 20 mg/ml tamoxifen (Sigma, T5648) dissolved in corn oil to induce deletion in all Tyrosinase expressing cells. 8-12 week old mice were injected daily with 100 µl tamoxifen for 5 consecutive days. Tumor development was measured weekly using calipers.

### Zebrafish Studies

Constructs *Pmitfa:EGFP:pA* and *Pmitfa:YAP-5SA:pA* were used in the miniCoopR assay as previously described (60). Briefly, *mitfa(lf)* mutant animals were bred, and single-cell stage embryos were injected with 25 pg of a single construct and 25 pg of Tol2 transposase mRNA. Successful Tol2-mediated integration of the construct into the genome rescued the *mitfa(lf)* phenotype, enabling melanocyte development. Melanocyte rescue was scored at 4-5 days of development, and rescued animals were grown to adulthood and monitored weekly for the presence of melanomas.

### Statistical analysis

All quantitative data are presented as mean +/-SEM, unless otherwise indicated. The number of samples (n) represents the number of biologic replicates, unless otherwise indicated. Prism 9 was used for all statistical analyses and for the creation of most graphs.

## Abbreviations

TCGA: The Cancer Genome Atlas
SKCM: skin cutaneous melanoma
MSKCC: Memorial Sloan Kettering Cancer Center
IB: Immunoblot
EV: empty vector

## RNAi Sequences

The siRNA’s used in this study are as follows:

Non-targeting #1 (Control siRNA) (Dharmacon):

UGGUUUACAUGUCGACUAA

LATS1 ON-TARGETplus SMARTpool (Dharmacon):

GGUGAAGUCUGUCUAGCAA; UAGCAUGGAUUUCAGUAAU

GGUAGUUCGUCUAUAUUAU; GAAUGGUACUGGACAAACU

LATS2 ON-TARGETplus SMARTpool (Dharmacon):

GCACGCAUUUUACGAAUUC; ACACUCACCUCGCCCAAUA;

AAUCAGAUAUUCCUUGUUG; GAAGUGAACCGGCAAAUGC

---

**Supplemental Figure 1.**
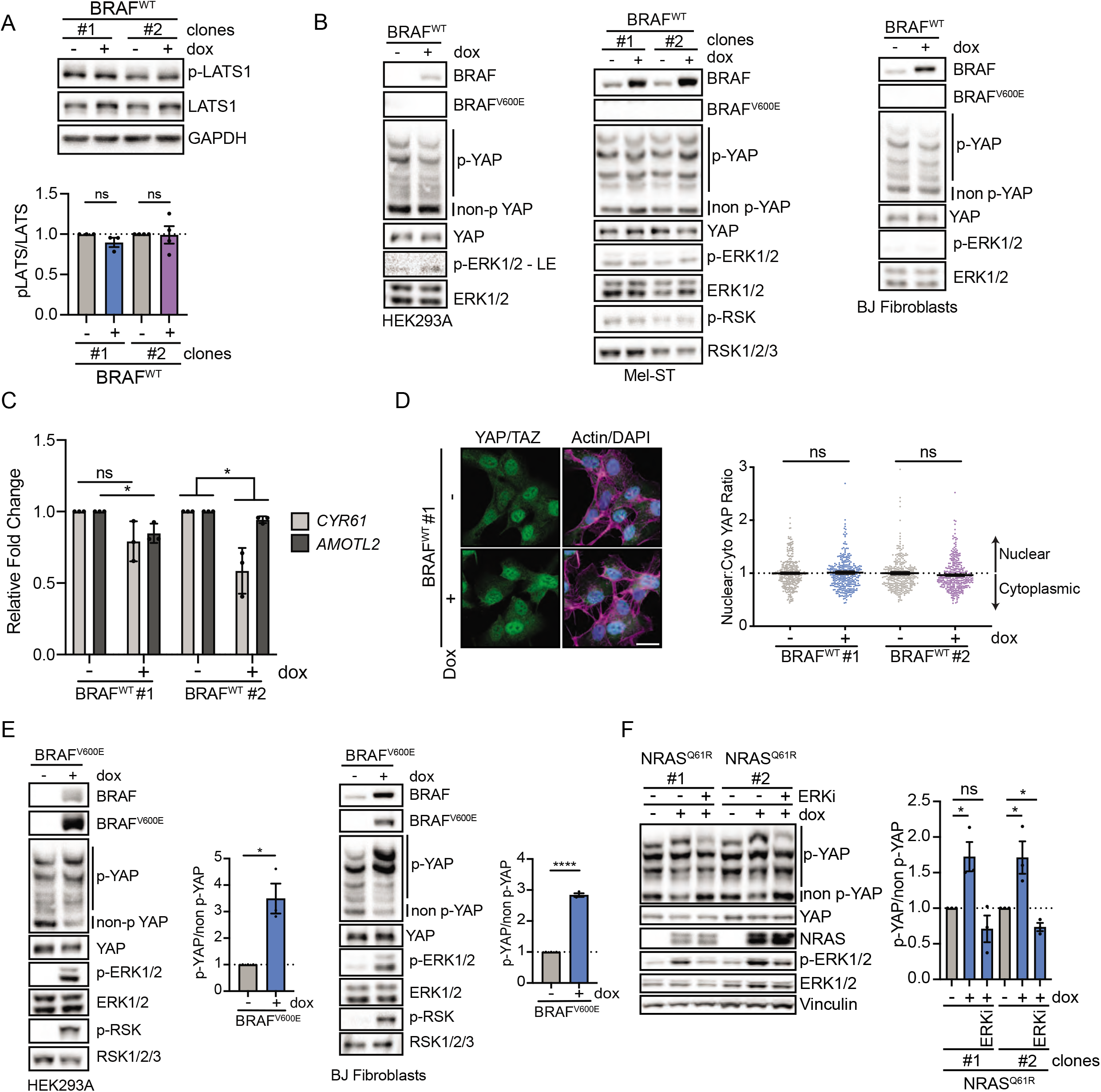
**(A)** IB of indicated *BRAF*^*V600E*^ dox-inducible Mel-ST clones cultured ± dox for 24 h (n ≥ 3 independent experiments, graph shows mean relative intensity ± SEM, two-tailed unpaired t test). **(B)** Representative IB of indicated *BRAF*^*WT*^ dox-inducible HEK293A, Mel-ST, and BJ Fibroblast cell lines (n = 3 independent experiments with similar results). **(C)** Relative expression of indicated genes in *BRAF*^*WT*^ Mel-ST clones cultured ± dox for 24 h (n = 3 independent experiments, graphs show mean ± SEM, two-tailed unpaired t test). **(D)** Left, representative immunofluorescence image of indicated *BRAF*^*WT*^ Mel-ST clone stained for YAP/TAZ or Merge of DAPI (blue), YAP/TAZ (green), and Phalloidin (magenta); Right, quantification of nuclear to cytoplasmic ratio of mean YAP/TAZ fluorescence (n > 300 cells from three independent experiments, graphs show mean ± SEM, scale bar 25 µm, two-tailed Mann-Whitney test). **(E)** Representative IB of indicated *BRAF*^*V600E*^ dox-inducible HEK293A and BJ cell lines with intensity quantification ratio of YAP phos-tags (n = 3 independent experiments, graphs show mean ± SEM, two-tailed unpaired t test). **(F)** Left, representative IB of *NRAS*^*Q61R*^ dox-inducible Mel-ST clones cultured ± dox for 24 h; Right, intensity quantification ratio of YAP phos-tag (n = 3 independent experiments, graphs show mean ± SEM, two-tailed unpaired t test). ns = non-significant, \**P* < 0.05,\*\**P* < 0.01, \*\*\**P* < 0.001, \*\*\*\**P* < 0.0001

**Supplemental Figure 2.**
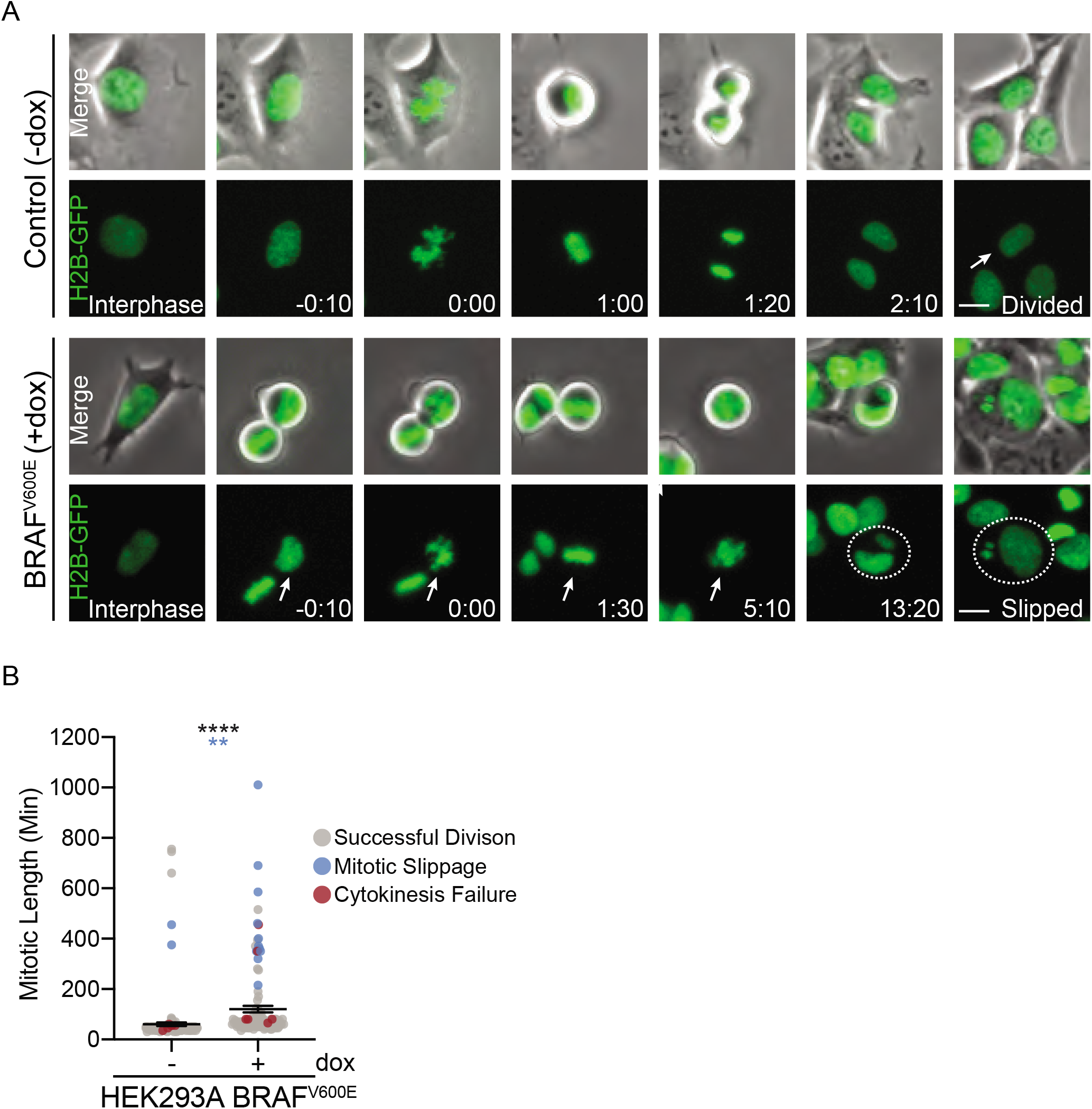
**(A)** Representative still fluorescence and bright-field images from a live-cell video of H2B-GFP (green) expressing *BRAF*^*V600E*^ dox-inducible HEK293A cells cultured ± dox (scale bar 25 μm, hh:mm). **(B)** Plot of mitotic length and fate of individually tracked mitoses from (A) (n > 120 mitoses per condition from two independent experiments, dots represent individually tracked mitoses, black stars represent mitotic length significance, two-tailed unpaired t test, blue stars represent significance for difference in frequency of whole-genome doubling events, two-sided Fisher’s exact test). \*\**P* < 0.01, \*\*\**P* < 0.001, ***** < 0.0001

**Supplemental Figure 3.**
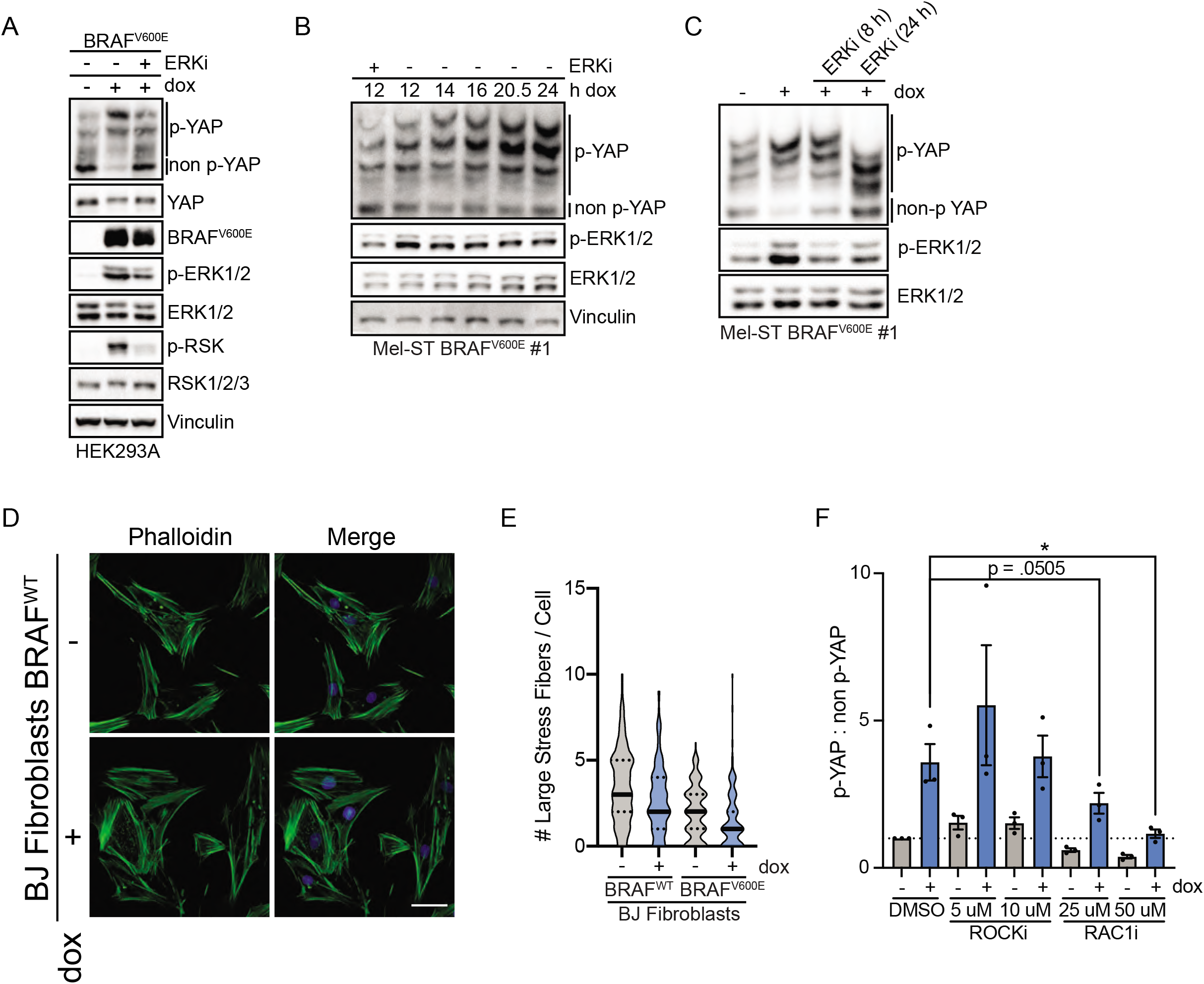
**(A)** IB of *BRAF*^*v600E*^ dox-inducible HEK293A cells treated with indicated drugs for 24 h (n = 3 independent experiments all with similar results) **(B)** IB of *BRAF*^*V600E*^ dox-inducible Mel-ST cell line treated with dox or ERKi for indicated times (n = 1 independent experiment) **(C)** Representative IB of *BRAF*^*V600E*^ dox-inducible Mel-ST cell line treated ± dox for 24 h followed by treatment of an ERK inhibitor for indicated time (n = 2 independent experiments with similar results). **(D)** Representative maximum intensity projections of confocal z-stacks of *BRAF*^*WT*^ dox-inducible BJ Fibroblasts cells stained for phalloidin and DAPI (n = 2 independent experiments with similar results, scale bar 25 μm). **(E)** Quantification of large stress fibers per cell in Figure 4D and S4D (n > 140 cells per condition from 2 independent experiments). **(F)** Intensity quantification ratio of YAP phos-tag from Figure 4E (n = 3 independent experiments, graph shows mean ± SEM, two-tailed unpaired t test). \**P* < 0.05

**Supplemental Figure 4.**
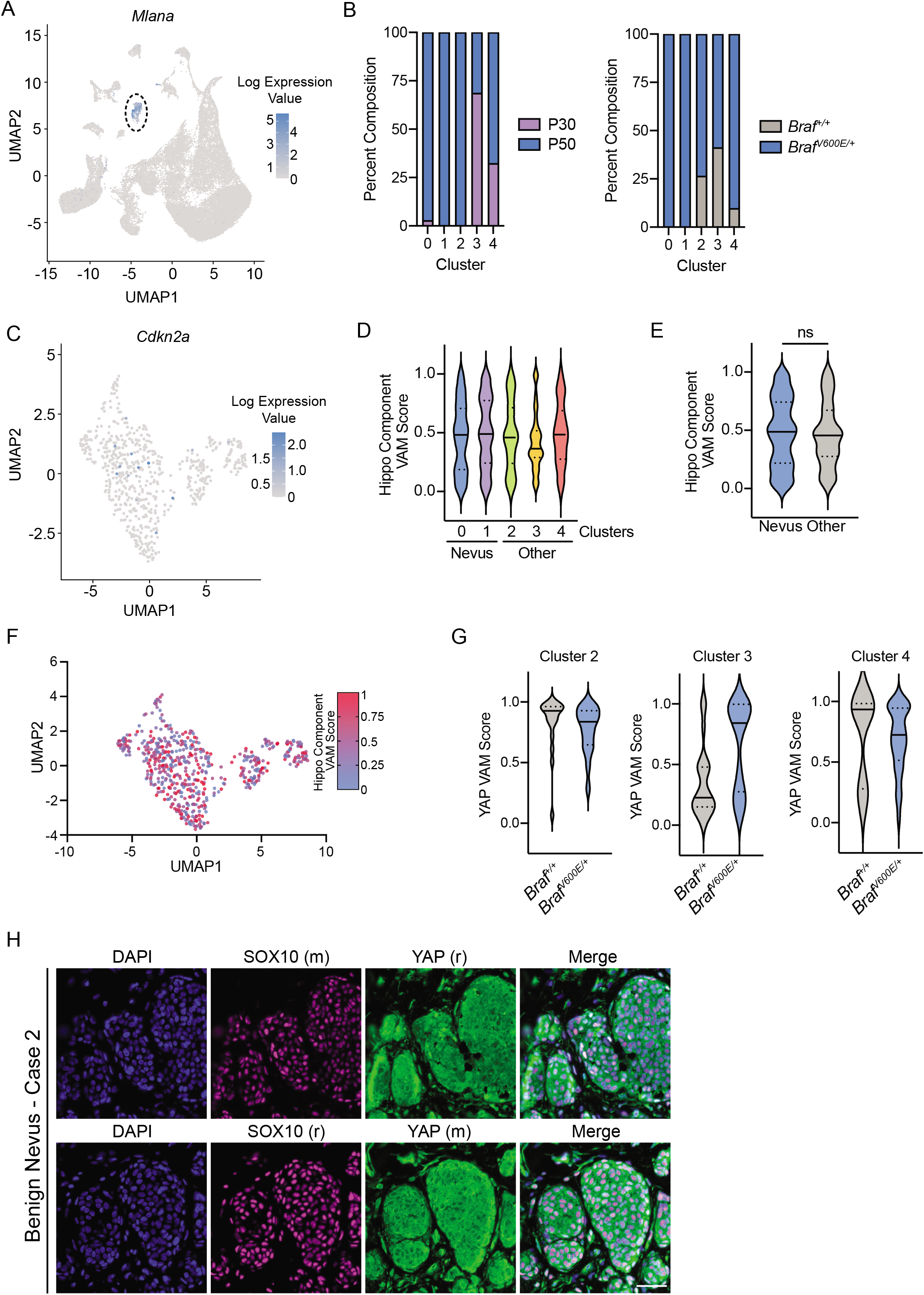
**(A)** UMAP of all single-cells from GSE154679 displaying relative expression of melanocyte marker *Mlana*. **(B)** Bar graphs of melanocyte subcluster composition dependent upon animal age (left) or genotype (right). **(C)** UMAP of *Cdkn2a* log expression across melanocyte subclusters (n = 589). **(D)** Hippo component VAM score plotted by melanocyte subcluster (n = 589) **(E)** Hippo component VAM score comparing nevus (clusters 0, 1) and other melanocytes (clusters 2, 3, 4) (nevus n = 408, other n = 181, two-tailed Mann-Whitney test). **(F)** UMAP of melanocytes colored by indicated gradient dependent upon Hippo Component VAM score (n = 589). **(G)** YAP VAM score according to indicated melanocytic subcluster and genotype. **(H)** Representative immunofluorescence staining of indicated proteins in another benign nevus case with two different sets of antibodies, (r) = rabbit, (m) = mouse, DAPI (blue), YAP (green), SOX10 (magenta), scale bar 50 µm. ns = non-significant

**Supplemental Figure 5:**
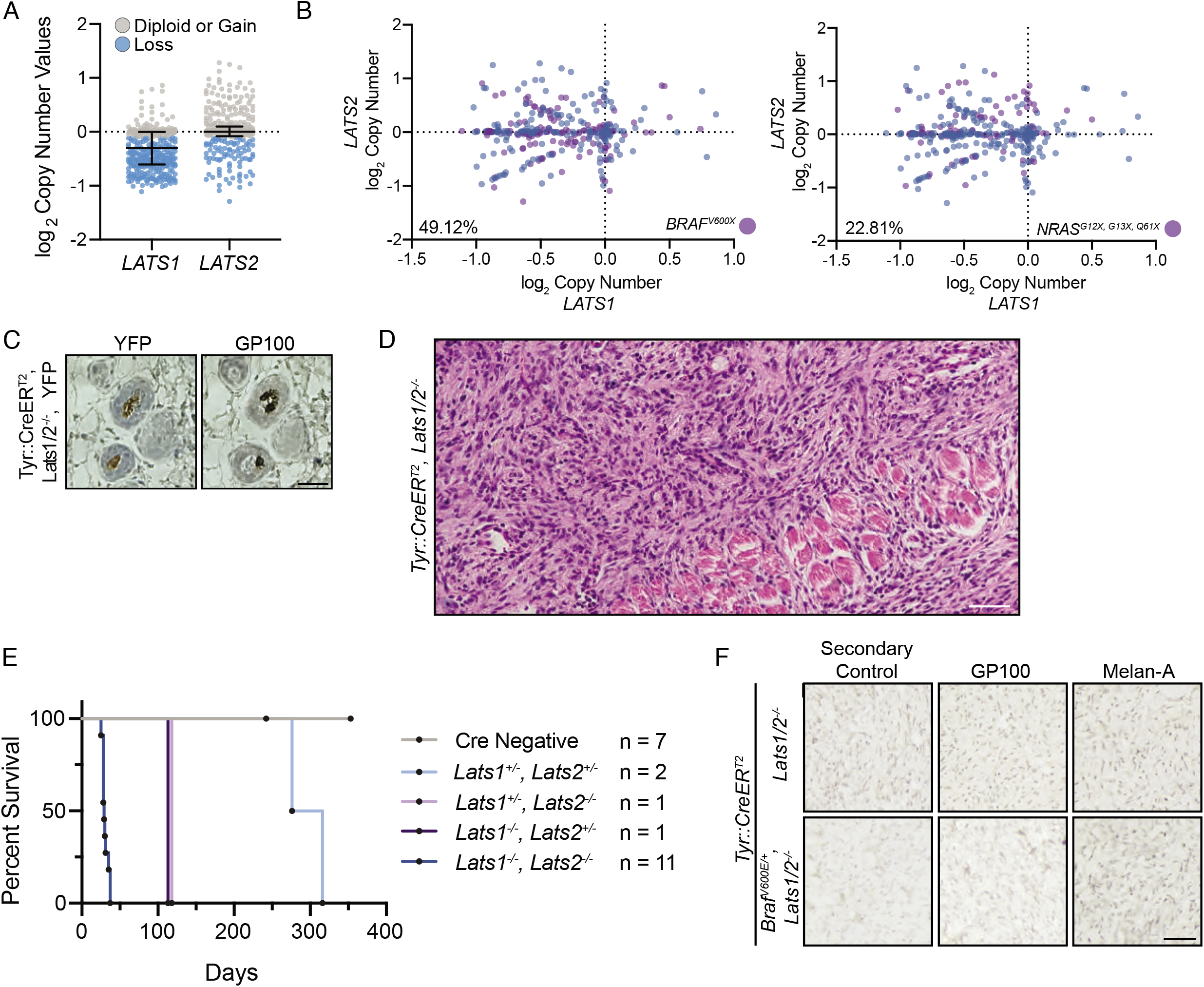
**(A)** Plot of log_2_ copy number Values from TCGA-SKCM for indicated genes. **(B)** Seme data as (A) with purple circles representing tumors with indicated mutations; bottom left percent is frequency of *LATS1/2* co-heterozygous loss that occurs with indicated mutations. **(C)** Representative IHC of indicated antigens, scale bar 40 μm (n ≥3 independent mice with similar results). **(D)** Hematoxylin and eosin staining of an early timepoint flank *Lats1/2*^*-/-*^tumor invading into underlying tissue, scale bar 50 μm. **(E)** Time from IP injection of tamoxifen to a study endpoint of mice with indicated genotypes. **(F)** Representative IHC of indicated antigens including secondary antibody only control, scale bar 40 um (n > 3 *Lats1/2*^*-/-*^n = 2 independent *Braf, Latsl/2*^*-/-*^mice with similar results).

**Supplemental Figure 6:**
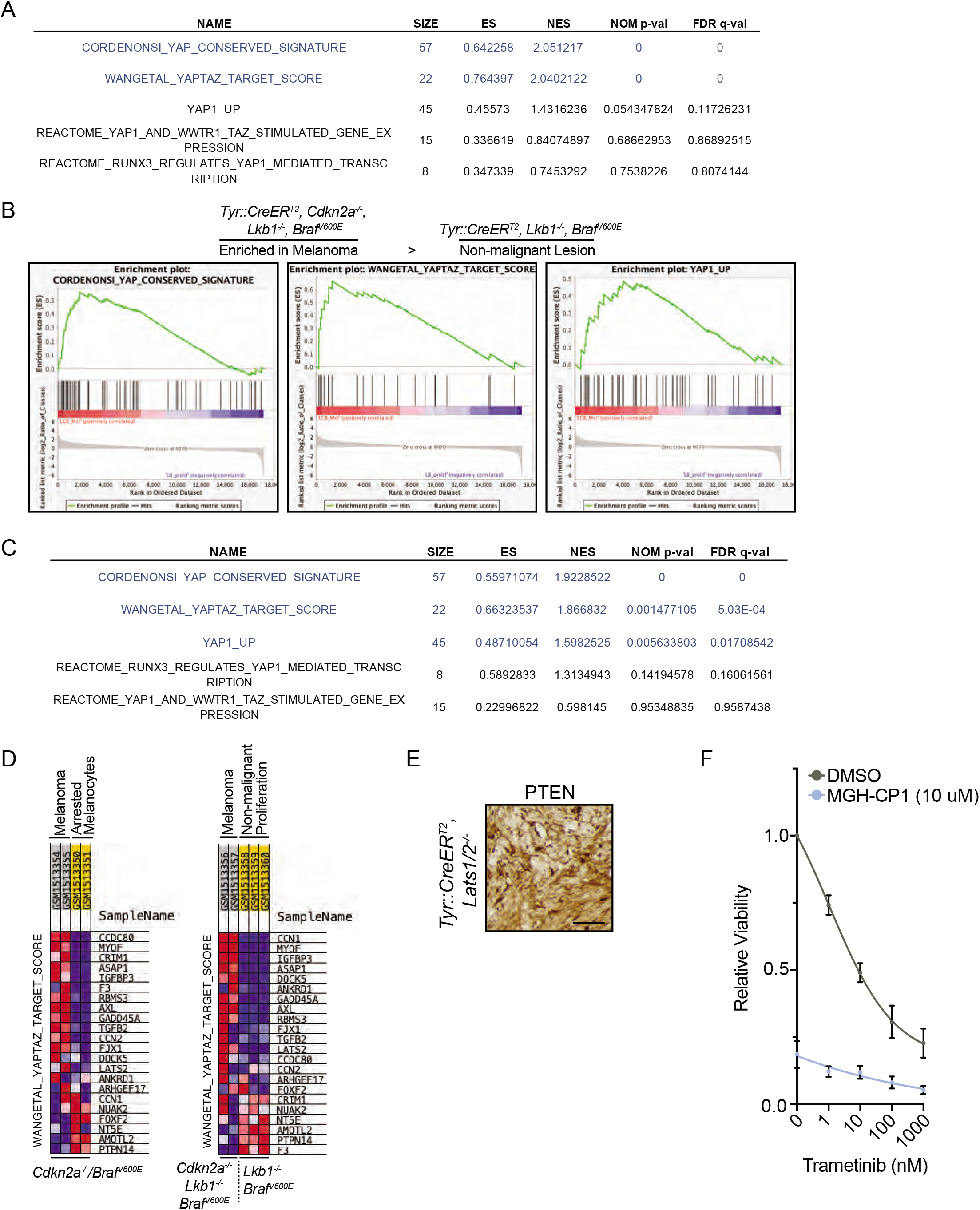
**(A)** Table of GSEA results from Figure 6F. **(B)** GSEA performed on GSE61750 comparing enrichment of YAP/TAZ signatures in melanoma to benign, proliferating melanocytes in indicated genotypes (see Figure S6D). **(C)** Table of GSEA results from Figure S6B. **(D)** GSEA heatmap results of gene expression from Figure 6F and S6B (red indicates higher expression, blue lower). **(E)** Representative IHC of PTEN, scale bar 40 µm (n = 2 independent mice with similar results). **(F)** Relative viability dose curve of indicated drugs in D4M.3A cells (n = 3 independent experiments in technical duplicate, graph shows mean ± SEM).

**Supplemental Table 1.**

| **YAP/TAZ Gene Set** | **Derived from:** |
| --- | --- |
| AGFG2 | Cordenonsi |
| AMOTL2 | Cordenonsi / Wang et al. 2018 |
| ANKRD1 | Cordenonsi / Wang et al. 2018 |
| ARHGEF17 | Wang et al. 2018 |
| ASAP1 | Cordenonsi / Wang et al. 2018 |
| AXL | Cordenonsi / Wang et al. 2018 |
| BICC1 | Cordenonsi |
| BIRC5 | Cordenonsi |
| CAVIN2 | Cordenonsi |
| CCDC80 | Wang et al. 2018 |
| CDC20 | Cordenonsi |
| CDKN2C | Cordenonsi |
| CENPF | Cordenonsi |
| COL4A3 | Cordenonsi |
| CRIM1 | Cordenonsi / Wang et al. 2018 |
| CCN2 (CTGF) | Cordenonsi / Wang et al. 2018 |
| CCN1 (CYR61) | Cordenonsi / Wang et al. 2018 |
| DAB2 | Cordenonsi |
| DDAH1 | Cordenonsi |
| DLC1 | Cordenonsi |
| DOCK5 | Wang et al. 2018 |
| DUSP1 | Cordenonsi |
| DUT | Cordenonsi |
| ECT2 | Cordenonsi |
| EMP2 | Cordenonsi |
| ETV5 | Cordenonsi |
| F3 | Wang et al. 2018 |
| FGF2 | Cordenonsi |
| FJX1 | Wang et al. 2018 |
| FLNA | Cordenonsi |
| FOXF2 | Wang et al. 2018 |
| FSCN1 | Cordenonsi |
| FSTL1 | Cordenonsi |
| GADD45A | Wang et al. 2018 |
| GADD45B | Cordenonsi |
| GAS6 | Cordenonsi |
| GGH | Cordenonsi |
| GLS | Cordenonsi |
| HEXB | Cordenonsi |
| HMMR | Cordenonsi |
| IGFBP3 | Wang et al. 2018 |
| ITGB2 | Cordenonsi |
| ITGB5 | Cordenonsi |
| LATS2 | Wang et al. 2018 |
| LHFPL6 | Cordenonsi |
| MARCKS | Cordenonsi |
| MDFIC | Cordenonsi |
| MYOF | Wang et al. 2018 |
| NDRG1 | Cordenonsi |
| NT5E | Wang et al. 2018 |
| NUAK2 | Wang et al. 2018 |
| PDLIM2 | Cordenonsi |
| PHGDH | Cordenonsi |
| PMP22 | Cordenonsi |
| PTPN14 | Wang et al. 2018 |
| RBMS3 | Wang et al. 2018 |
| SCHIP1 | Cordenonsi |
| SERPINE1 | Cordenonsi |
| SGK1 | Cordenonsi |
| SH2D4A | Cordenonsi |
| SHCBP1 | Cordenonsi |
| SLIT2 | Cordenonsi |
| STMN1 | Cordenonsi |
| TGFB2 | Cordenonsi / Wang et al. 2018 |
| TGM2 | Cordenonsi |
| THBS1 | Cordenonsi |
| TK1 | Cordenonsi |
| TNNT2 | Cordenonsi |
| TNS1 | Cordenonsi |
| TOP2A | Cordenonsi |
| TSPAN3 | Cordenonsi |

**Supplemental Table 2.**
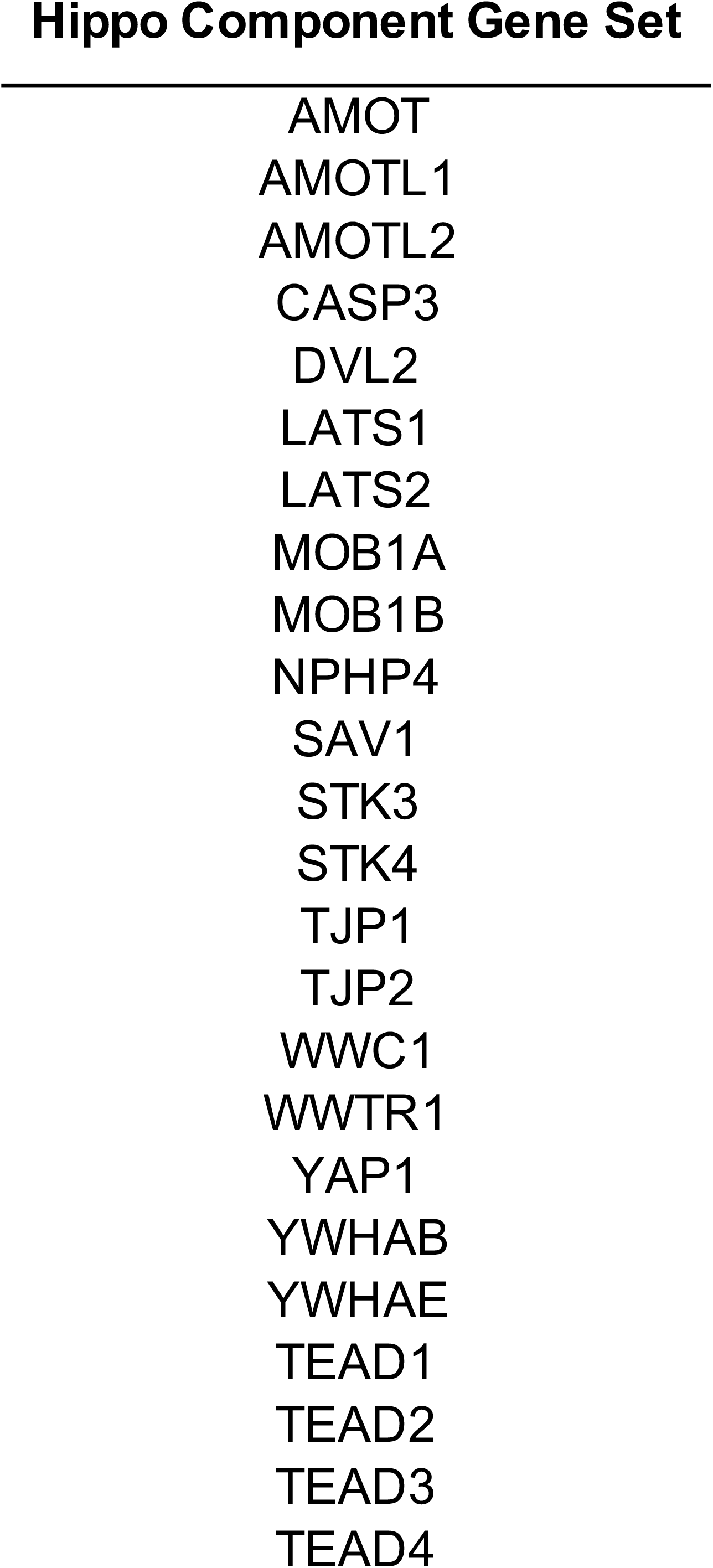

